# Comparative fitness of reemerging St. Louis encephalitis virus in vertebrate and mosquito cells, *Culex tarsalis* and *Culex quinquefasciatus* mosquitoes, and mice

**DOI:** 10.64898/2025.12.16.694562

**Authors:** M. Arturo Flores Rodriguez, Hongwei Liu, Erik Turner, Elias Im, Rochelle Leung, Sunny An, Christopher M. Barker, Lark L. Coffey

## Abstract

The human pathogenic orthoflavivirus St. Louis encephalitis virus (SLEV) reemerged in the western United States in 2015 after more than a decade of absence and has since expanded throughout California with sustained interannual transmission. This shift from the historically sporadic pattern of SLEV activity before 2003 raises the question of whether contemporary strains differ in fitness from earlier strains. To assess whether reemerging SLEV possess enhanced infectivity or transmissibility, we compared five contemporary genotype III strains from California (2016–2023) with a historical genotype V strain from 2003. Growth kinetics were evaluated in mammalian, duck, and mosquito cells; vector competence was assessed in laboratory colonies of *Culex tarsalis* and *Culex quinquefasciatus* vectors; and viremia profiles were measured in Collaborative Cross recombinant intercross mice. Some genotype III strains produced higher titers than the historical genotype V strain in avian and mosquito but not mammalian cells. Several genotype III strains infected and transmitted SLEV RNA more efficiently than the historical strain in both mosquito species, although no temporal trend in fitness was observed. SLEV fitness was comparable or greater in *Culex quinquefasciatus* than in *Culex tarsalis*. Sequencing identified no shared amino acid substitutions associated with vector infection phenotypes. Although genotype III strains exhibited a delayed peak relative to the historical strain, murine viremia levels were comparable across strains. These findings show some contemporary strains exhibit equal or greater fitness than the historical strain, which may contribute to SLEV persistence and spread in California, underscoring the need for continued surveillance and targeted vector control.

**IMPORTANCE:** St. Louis encephalitis virus (SLEV) reemerged in California in 2015 after more than a decade of absence and has since established sustained transmission and expanded geographically. The factors underlying this reemergence remain poorly understood. By comparing contemporary genotype III SLEV strains with a historical genotype V strain, we found that several contemporary strains exhibit equal or greater fitness compared to the historical strain in avian and mosquito cells and are transmitted more efficiently by the two principal California vector species, *Cx. tarsalis* and *Cx. quinquefasciatus*. We also demonstrate that *Cx. quinquefasciatus* can transmit infectious genotype III SLEV, supporting its role in SLEV maintenance and spread. Despite differences in mosquito infection and transmission, we found no evidence that fitness in mosquito vectors or mice has continued to rise among strains detected more recently, suggesting that enhanced transmission is not driven by ongoing directional adaptation. These findings indicate that contemporary genotype III SLEV strains possess transmission competence in mosquito vectors that may have contributed successful reestablishment and persistence of SLEV California. Improved understanding of the characteristics of reemerging SLEV strains can inform surveillance, risk assessment, and vector control efforts aimed at reducing human exposure to prevent disease caused by SLEV.

## INTRODUCTION

St. Louis encephalitis virus (SLEV, *Flaviviridae, Orthoflavivirus louisense*) circulates in an enzootic cycle between birds and mosquitoes in the Americas and can spill over into humans, causing febrile illness and sometimes encephalitis. SLEV produced periodic outbreaks in the United States (US) beginning in the 1930s and was a major cause of encephalitis during the 1970s (1, 2), including in California from the 1940s through the 1990s (3, 4). Because no specific treatment or vaccine for SLEV exists, mosquito control remains the primary public health tool for limiting SLEV transmission. In California, public health agencies monitor SLEV activity using human cases, sentinel chicken seroconversions, mosquito surveillance, and the distribution and abundance of mosquito vectors. Mosquito surveillance showed near annual SLEV activity from 1970 until 2003 (3), the year the related flavivirus West Nile virus (WNV) invaded the state (5). Starting the next year (2004) until 2015, SLEV was not detected in California mosquitoes. The reasons for the disappearance of SLEV are not fully understood, but sterilizing cross-protection by WNV (6) and high levels of WNV herd immunity in shared avian reservoirs (7) may have quenched SLEV activity after WNV invaded. In 2015, a SLEV outbreak occurred in Phoenix, Arizona, concurrent with a WNV epidemic (8). That same year, SLEV was detected again in California and subsequently spread northward from southern California into the Central Valley and farther north from 2016-2025 (9) into 22 of 58 (38%) of California counties. Since 2015, SLEV-positive mosquito pools have included *Cx. stigmatosma, Cx. pipiens, Cx. tarsalis*, and *Cx. quinquefasciatus* (10), the latter three of which were previously identified as the primary vectors of both WNV and historic SLEV in California (11–14). Between 2015-2025, 95% of SLEV positive pools in California were *Cx. tarsalis* or *Cx. quinquefasciatus* (10). The geographic expansion, increasing number of human cases most years, and detection of SLEV positive mosquito pools since 2015, shows SLEV is becoming increasingly prevalent in California (10). This pattern contrasts with historical pre-2003 SLEV activity, during which the virus was typically locally introduced, amplified for a 1-2 year period, and then disappeared (4, 15–17). Moreover, although SLEV and WNV now co-circulate in many of the same California counties, SLEV remains largely absent from WNV hotspots in the north and south and is instead more common in the Central and Coachella Valleys (9). The interannual persistence and geographic spread of reintroduced SLEV since 2015 raises the question of whether its reestablishment is mediated by increased fitness, defined here as infectivity and transmissibility in mammalian, duck, and mosquito cells, colonies of the two primary California mosquito vector species *Cx. tarsalis* and *Cx. quinquefasciatus*, and mice.

Our analyses comparing SLEV genomes in the western US from 2015-2018 with publicly available sequences show that these strains are most closely related to SLEV associated with an outbreak in Argentina in 2005 (18) and to a 2014 isolate from Argentine mosquitoes (19), supporting a South American introduction (20, 21). SLEV circulating in California and the western US from 2014-2018 forms at least 4 geographically distinct clusters (a-d), all belonging to genotype III (21). Subsequent SLEV phylogenetic studies conducted after 2018 indicate that multiple SLEV importations have occurred in California, Arizona, and other western states, most genomes belong to genotype III (22); Texas also reports genotype II (23). To our knowledge, 2014 was the first time genotype III SLEV was detected in the US, although Bayesian phylogenetics suggest it was present in the US at least a year earlier, in 2013 (23). Despite these genetic insights, no phenotypic studies in cells, mosquito vectors, or mice have characterized infection or transmission dynamics of reemerging genotype III SLEV, nor compared post-2015 strains with pre-2004 SLEV (e.g. strains from 2003, which belong to genotype V that is not the direct ancestor of genotype III), that historically circulated in California and have been extensively evaluated in mosquito and avian hosts (7, 16, 24–29). We therefore sought to compare the relative fitness of reemerging post-2015 genotype III SLEV with pre-2003 genotype V, where genotype V was last genotype detected in California prior to the 11-year absence of SLEV activity in mosquitoes. For context, WNV, which shares many avian reservoirs and *Cx.* vector species with SLEV, has evolved to increase transmission competence in *Cx.* spp. across the US since 2002 (30). Work in California further showed that WNV strains from later outbreaks (2004–2012) outcompete a 2003 strain in house finches but not in *Cx. tarsalis* (31, 32), suggesting that enhanced avian rather than mosquito fitness likely contributed to WNV amplification to outbreak levels during a decade of circulation in California. Although WNV evolution in both vectors and avian hosts across the US and in California is well documented, comparable studies for SLEV are lacking, leaving unanswered whether changes in viral fitness contributed to SLEV reemergence in California after 2015. To address this gap, the goal of this study was to evaluate the extent to which SLEV reemergence may be driven by altered infectivity and transmissibility in mammalian, duck, and mosquito cells, two established colonies of primary California mosquito vectors, and Collaborative Cross recombinant intercross mice (RIX). The collaborative cross RIX mice were chosen as an immunocompetent mice model of flavivirus disease. Prior studies show they are susceptible to infection with WNV (33, 34) and tick-borne Powassan virus (35) and develop detectable viremias within a week after inoculation. Furthermore, in contrast to established murine SLEV models employing immunocompromised mice (36) and intracranial inoculation (37), we sought to use mouse model that supports infection after peripheral inoculation. We compared fitness of one historic SLEV strain from 2003 that is in genotype V and five SLEV genotype III strains from 2016-2023 in each of these systems.

## MATERIALS AND METHODS

### Biosafety

All work with SLEV was performed in a biosafety level 3 laboratory at the University of California, Davis, under approved Biological Use Authorization #R1863.

### Cell culture

African green monkey kidney (Vero, ATCC CCL-81) cells were used for SLEV isolation from mosquito pools, growth curves, and virus titrations from growth curves, bloodmeals, and mosquito saliva. Vero cells were maintained in Dulbecco’s Modified Eagle’s Medium (DMEM) supplemented with 5% heat-inactivated fetal bovine serum (FBS), 100 U/mL penicillin and 100 μg/mL streptomycin at 37°C with 5% CO_2_ in a humidity-controlled incubator. Primary *Cx. Tarsalis* (CxTr) were used for growth curve experiments. CxTr cells were a gift from Dr. Dane Jasperson at the United States Department of Agriculture. CxTr cells were maintained at 27°C in the dark in modified Schneider’s insect medium prepared from Schneider’s medium supplemented with sodium bicarbonate, L-glutamine, L-asparagine, reduced glutathione, and 10% heat-inactivated FBS. Media additions were filter sterilized prior to use. CxTr cells were maintained as closed cultures without supplemental CO_2_ using vented-cap flasks. Duck embryonic fibroblast (DEF) cells (ATCC CCL-141) were used for growth curve experiments. DEF were maintained in DEF cell culture medium consisting of DMEM supplemented with 10% heat-inactivated FBS and 1% penicillin-streptomycin at 37°C with 5% CO_2_ in a humidity-controlled incubator.

### SLEV activity in California mosquitoes

The total annual number of SLEV positive mosquito pools and SLEV detections in each mosquito species in California were sourced from the California Department of Public Health Vector Borne Disease Section Annual Reports from 2015-2024 (10). For the 2016 report, detailed data on the numbers of SLEV detections in different mosquito species were not provided; this data was instead derived from weekly bulletins from 2016 (38). The 2025 data was obtained from the WestNile.ca.gov website (39) and is current as of December 8, 2025.

### SLEV isolation from mosquito pools

This study focuses on contemporary SLEV from the period 2016-2023 and utilizes five genotype III SLEV strains isolated from California locations with ongoing mosquito pool activity compared to a historical genotype V strain from 2003. Pools of *Cx. tarsalis* and *Cx. quinquefasciatus* collected in California (**Table 1**) and confirmed SLEV positive by an established qRT-PCR method (40) were obtained through routine arbovirus surveillance conducted by mosquito abatement districts in California and the Davis Arbovirus Research and Training laboratory at the University of California, Davis. SLEV positive mosquito pools were centrifuged at 4,000 g for 5 minutes to pellet debris and then filtered through a 0.22 μM filter (Millex GP, Millipore, Billerica, MA, USA). The filtrate was inoculated onto confluent T25 Vero cell monolayers and incubated for 4 days. Virus recovered from culture supernatants was subsequently passaged 2-4 additional times on Vero cells to generate working stocks, which were titrated in Vero cells and stored at -70°C in aliquots until use. The genotype V SLEV strain isolated in 2003, representing the lineage historically circulating in California prior to 2004, was included as a comparator; this strain had been isolated previously and was re-sequenced since 2 additional Vero cell passages were performed to generate a working stock. The genotype III strains share ≍99% nucleotide and amino acid identity genome-wide with one another and ≍93% nucleotide and ≍98% amino acid identity with the 2003 genotype V strain (**Table 2**).

**Table 1.** SLEV strains used.

| Name | Strain identifier | Location | Year | Mosquito Source | Passage history | GenBank Accession number | Genotype |
| --- | --- | --- | --- | --- | --- | --- | --- |
| 2003_V | IMP115 | Imperial County, CA | 2003 | *Cx. tarsalis* | Vero P3 | PX678850 | V |
| 2016_IIId | KERN345 | Kern County, CA | 2016 | *Cx. quinquefasciatus* | Vero P4 | MN233315 | IIId |
| 2017_IIIa | IMPR570 | Imperial County, CA | 2017 | *Cx. tarsalis* | Vero P3 | MN233313 | IIIa |
| 2022_IIId | FRWS339 | Fresno West Side, CA | 2022 | *Cx tarsalis* | Vero P2 | PX678851 | IIId |
| 2023_IIIa | COAV3764 | Coachella Valley, CA | 2023 | *Cx tarsalis* | Vero P2 | PX678852 | IIIa |
| 2023_IIId | MERC702 | Merced, CA | 2023 | *Cx tarsalis* | Vero P2 | PX678853 | IIId |

**Table 2:** Genome-wide nucleotide / amino acid identity between SLEV strains used.

|  | 2003_V | 2016_IIId | 2017_IIIa | 2022_IIId | 2023_IIIa | 2023_IIId |
| --- | --- | --- | --- | --- | --- | --- |
| 2003_V | 100.0 / 100.0 | 93.04 / 98.69 | 93.03 / 98.75 | 93.15 / 98.72 | 93.07 / 98.72 | 93.16 / 98.69 |
| 2016_IIId | 93.04 / 98.69 | 100.0 / 100.0 | 99.66 / 99.77 | 99.61 / 99.74 | 99.50 / 99.71 | 99.38 / 99.65 |
| 2017_IIIa | 93.03 / 98.75 | 99.66 / 99.77 | 100.0 / 100.0 | 99.45 / 99.74 | 99.35 / 99.71 | 99.57 / 99.77 |
| 2022_IIId | 93.15 / 98.72 | 99.61 / 99.74 | 99.45 / 99.74 | 100.0 / 100.0 | 99.79 / 99.80 | 99.20 / 99.68 |
| 2023_IIIa | 93.07 / 98.72 | 99.50 / 99.71 | 99.35 / 99.71 | 99.79 / 99.80 | 100.0 / 100.0 | 99.10 / 99.65 |
| 2023_IIId | 93.16 / 98.69 | 99.38 / 99.65 | 99.57 / 99.77 | 99.20 / 99.68 | 99.10 / 99.65 | 100.0 / 100.0 |

### SLEV growth curves

Vero, DEF, and CxTr cells were seeded in 24-well plates one day prior to SLEV inoculation to achieve approximately 80% confluence at the time of inoculation. Triplicate wells of cells were inoculated with SLEV strains at a multiplicity of infection (MOI) of 0.1 in 100 µL per well, which was incubated for 1 hour at 37°C with rocking every 10 minutes. After the incubation, cells were washed twice with DPBS and overlaid with 1 mL of growth medium. Supernatants were collected at 0, 1, 2, 3, 4, 5, 6, and 7 days post inoculation (dpi) by transferring 20 µL from each well into 180 µL of viral diluent consisting of DPBS with 1% FBS, followed by immediate storage at −80°C.

### SLEV titrations and infectious virus detection in saliva

Titers of SLEV stocks, growth curve samples, and bloodmeals were quantified by plaque assays using Vero cells. The presence of infectious virus was qualitatively assessed in a subset of randomly selected saliva samples that had detectable SLEV RNA. For SLEV stocks, growth curve samples, and bloodmeals, serial tenfold dilutions of virus were inoculated onto confluent monolayers in 6-well plates. For saliva, confluent monolayers in 12-well plates were inoculated with 50 µl of mosquito saliva in DMEM. Inocula were adsorbed to cells for 1 hour at 37°C with 5% CO_2_ with gentle rocking every 15 minutes. Following adsorption, an agarose-based overlay containing nutrient medium, 0.5% agarose and 3% bicarbonate (Sigma Aldrich, St. Louis, MO, USA) was added. After 5 days of incubation at 37°C and 5% CO_2_, a second overlay containing additional 3% neutral red (Sigma Aldrich, St. Louis, MO, USA) was applied. On 7 dpi plaques were visualized and, for stocks, growth curve samples, and bloodmeals, enumerated to determine the titer in log_10_ plaque forming units (PFU)/mL. Saliva samples were tested once on one well due to volume constraints. Saliva are reported as positive if at least 1 plaque was detected. Each stock or bloodmeal titration was performed in duplicate, and the mean value is reported. SLEV titers reported from growth curves represent the mean titer from triplicate wells, where virus in each well was measured in one titration.

### Mosquitoes

Laboratory colonies of *Cx. tarsalis* and *Cx. quinquefasciatus* were used. The *Cx. tarsalis* colony originated from collections in 2002 at the Kern National Wildlife Refuge in Kern County, California. The *Cx. quinquefasciatus* colony (CQ1) originated from collections in Merced County, California in the 1950s (41). For simplicity, we refer to the CQ1 colony as *Cx. quinquefasciatus* but given the geographic origin of the colony within the zone of hybridization with *Cx. pipiens* (42), the CQ1 colony may represent an admixture of the two species. Both colonies have been maintained continuously since establishment. Mosquitoes were reared under insectary conditions of 22°C, ∼30% relative humidity, and 12 hour:12 hour light:dark cycle. Larvae were maintained in 1 liter deionized water at 200-400 larvae per pan and fed 1 pinch of fish food (Tetra, Melle, Germany) every other day until pupation. Adults were housed in 32.5 x 32.5 x 32.5 cm cages (BugDorm, #4F3030, Megaview Science, Taiwan) with constant access to 10% sucrose. Adults aged 3-7 days were used for vector competence experiments.

### Vector competence

A total of five genotype III SLEV strains were assessed; three were tested in *Cx. tarsalis* and all were tested in *Cx. quinquefasciatus*. The historical 2003 genotype V strain was included for comparison in both species. Approximately 200 mixed-sex mosquitoes were aspirated from colony cages 1 day prior to bloodfeeding and transferred into 946 cm^3^-sized plastic containers with mesh lids and access to 10% sucrose. SLEV stocks were diluted in heparinized sheep blood (HemoStat Laboratories, Dixon, CA, USA) to target doses of 3, 5, and 6.7 log_10_ PFU/ml. The 3 and 5 log_10_ PFU/ml doses are within the range of avian reservoir viremias, whereas the 6.7 log_10_ PFU/ml dose exceeds avian viremias (43). Because infectious arbovirus titers can be affected by storage and freeze–thaw conditions (44), our use of previously frozen virus stocks may have reduced the effective infectious dose relative to the measured target titer, potentially bringing the highest dose closer to physiologically relevant avian viremia levels. Bloodmeals were offered for 60 minutes using a collagen membrane mounted on a Hemotek member feeder (Hemotek Ltd, Blackburn, United Kingdom) maintained at 37°C. Fully engorged females, identified by visible blood in the abdomen, were anesthetized for 10 seconds with CO_2_, sorted into clean plastic containers with mesh lids at a density of 30-60 mosquitoes, and held at 28°C with 60-70% humidity and 12 hour:12 hour light:dark cycle for 13, 14, or 15 days, with constant access to 10% sucrose. A total of 6 experiments were conducted in which 2-4 SLEV strains were compared at matched bloodmeal titers. Each strain was tested in 1-4 experiments; the combined data from all experiments are shown.

At 13, 14, or 15 days post-bloodfeeding, mosquitoes were CO_2_-anesthetized and placed immobile on a chill table for dissection. Harvests were distributed across the three day period due to personnel capacity constraints. Legs and wings were removed prior to saliva collection, which was performed by inserting the proboscis into capillary tubes containing FBS for 20-30 minutes. Each capillary tube was placed in a 1.5 mL tube containing 100 µL DMEM and a 200 ul pipette tip was used to aspirate the FBS containing saliva from the capillary tube. Bodies and legs & wings were placed into 2 mL tubes (Thermo Fisher Scientific, Emeryville, CA) containing 250 µL DMEM and a 5-mm glass bead (Thermo Fisher Scientific, Emeryville, CA). Dissection tools were rinsed once in 70% ethanol between each sample to prevent cross-contamination. Tissues were homogenized at 30 Hz for 4 minutes in a TissueLyser (Retsch, Haan, Germany) and centrifuged at 10,000 g for 2 minutes, then stored at -70°C until processing.

### Mice

Collaborative Cross recombinant intercross (RIX) 024×023 mice were generated by the Systems Genetics Core Facility (SGCF) at the University of North Carolina. Triads of collaborative cross (CC) 024 and CC023 were used to generate F1 hybrids that were maintained at the highest health status barrier facility for breeding. The SGCF conducts yearly quality control genotyping, and individual ear punches from all mice used in this study were archived. All mouse procedures were approved by the University of California, Davis Institutional Animal Care and Use Committee under protocols #23308 and #24862 and were performed in accordance with approved procedures.

### Mouse infections

To compare replication kinetics of SLEV strains in blood, adult (8-12 week old) Collaborative Cross RIX 024×023 mice were subcutaneously inoculated in the right rear footpad with 100 plaque forming units (PFU) of one of six SLEV strains or mock inoculum (DPBS), each diluted in Vero cell growth medium. The inoculum volume was 10 µL. Each experimental group consisted of 12 mice (6 males and 6 females). Mice were monitored daily for weight change relative to day 0 weight from 1 to 4 dpi. Blood was collected via submandibular cheek bleed. To reduce repeated blood collection from individual animals, six mice per group were bled on 1 and 3 dpi, while a different set of six mice was bled on day 2 dpi. All mice were bled on day 4 dpi immediately prior to euthanasia. Four dpi was selected for euthanasia since the focus of these studies was the onset and early viremic period. Whole blood was immediately diluted 1:10 in DMEM and frozen at -80°C. Frozen diluted blood was thawed at room temperature prior to RNA extraction and viremia quantification by qRT-PCR.

### SLEV RNA detection, enumeration, and reporting

Tissues were thawed and viral RNA was extracted from mouse blood and portion of bodies and legs & wings using the manufacturer’s instructions for the MagMax Viral RNA Extraction Kit (ThermoFisher Scientific, Emeryville, CA, USA) or saliva using Trizol LS (ThermoFisher). For Trizol extractions, 400 µL TRIzol LS and 100 µL chloroform was added to 100 µL of saliva. The tube was mixed vigorously and then centrifuged for 5 minutes at 20,000 g at 4°C. The clear upper aqueous layer was transferred to a new 1.5 mL tube, and 200 µL of isopropanol was added. Then the tube was gently mixed by inverting 5-10 times followed by incubation for 30 minutes at -20°C. The sample was centrifuged for 15 minutes at 20,000 g at room temperature. The supernatant was discarded, and the sample was centrifuged for 30 seconds, and residual ethanol was removed using a 20 µL pipette. Total RNA isolated was suspended in 90 µL buffer following the MagMax manufacturer’s instructions (tissue samples) or 100 µL of nuclease-free water (saliva) from the Trizol method. SLEV RNA in mosquito bodies, legs & wings, and saliva was quantified by qRT-PCR using a TaqMan Fast Virus 1-Step Mastermix and a SLEV-specific primer set at 20 μM (**Table 3**) using the following reagents: 3.2 μl TE buffer, 1.8 μl of the primer and probe mix, 5 μl of Fast mix, and 10 μl of RNA. The cycling conditions were as follows: 50°C for 5 minutes, 95°C for 20 seconds, and 40 cycles of 95°C for 3 seconds followed by 60°C for 30 seconds. Cycle threshold (Ct) values were converted to RNA genome copies using standard curves generated from known SLEV RNA concentrations included on each qRT-PCR plate. At the consensus sequence level, all six SLEV strains are 100% identical across the forward primer and probe binding regions (**Supplemental Figure 1**). Both the forward primer and the probe exhibit a single nucleotide mismatch relative to all SLEV strains evaluated, with a second mismatch between the probe and 2003_V. Within the reverse primer-binding region, strains 2016_IIId and 2003_V each contain one nucleotide mismatch at distinct positions. The SLEV RNA of known concentration used to generate the standard curve was commercially procured from ATCC (Manassas, Virginia, Item# VR-3236SD) but the genomic sequence from which the primers and probe are designed is not reported by ATCC. Negative control samples lacking SLEV RNA were included in each run. Samples were assayed in technical duplicates or triplicates. Samples in which only one replicate produced a Ct < 40 were considered negative. The limit of detection (LOD) was defined as the mean Ct for the least concentrated standard curve point producing a Ct < 40 across all runs. Samples that did not yield a detectable Ct < 40 are not included in group means for titer measurements. SLEV RNA concentrations are reported as log_10_ SLEV genomes/ml mouse blood or total mosquito tissue or saliva sample calculated as geometric means and analyzed following log transformation. For vector competence studies, infection, dissemination, and transmission rates were calculated as the number of positive bodies, legs & wings or saliva samples, respectively, divided by the number of total samples of the same type that were tested, multiplied by 100. Using a common denominator allows direct comparison across stages of vector competence and, for transmission, provides an epidemiologically relevant estimate of the probability that a mosquito will become capable of transmitting virus after ingesting an infectious blood meal.

**Table 3:**
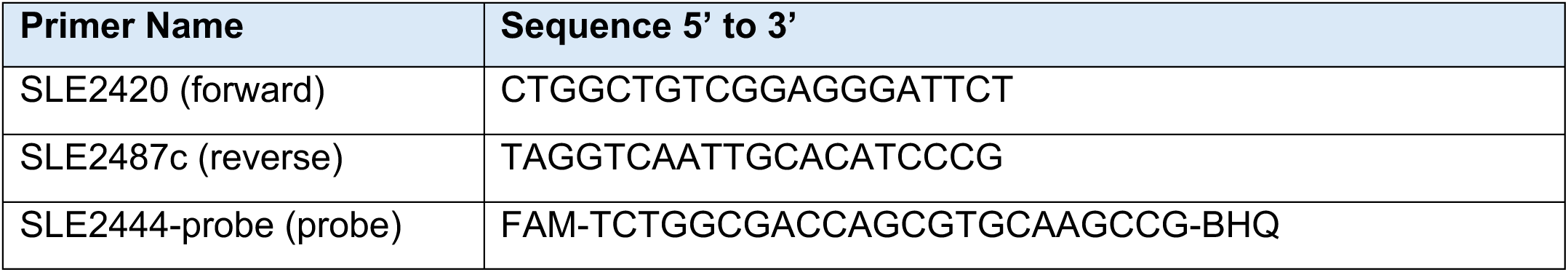
Primers used for SLEV RT-qPCR. FAM is fluorescein amidite and BHQ is black hole quencher.

### SLEV genetic comparisons

Alignments of SLEV genomes and primers and probe were generated to compare nucleotide and amino acid identity using the EMBL-EBI MAFFT multiple sequence alignment tool and CLUSTAL (45).

### SLEV positive mosquito pool mapping

Mosquito pool data were used with permission from (9) and mapped using ArcGIS Pro 3.1.

### Statistical Analyses

All statistical analyses were performed in GraphPad Prism version 10.3.1. Analyses of infectious SLEV titers (PFU) and SLEV RNA levels were conducted on log-transformed values. Growth kinetics in cells and viremia kinetics in mice were compared among SLEV strains using a mixed-effects model including strain, time, and their interaction, with cell replicate or mouse included as a random effect and 2-way ANOVA with Tukey’s multiple comparisons tests. Two-sided Fisher exact pairwise comparisons with Holm correction were used to compare rates of infection, dissemination, and transmission between mosquito species and SLEV strains in cohorts of mosquitoes presented the same bloodmeal titer. For RNA quantification, differences in mean RNA levels among SLEV strains in the same mosquito species were evaluated using ANOVA with Šídák’s multiple-comparisons test. Only adjusted p values ≤ 0.05 were considered statistically significant. The specific statistical tests used for each comparison are provided in the Results section.

### Data availability

All data supporting the findings of this study are available as Supplemental File 1.

## RESULTS

SLEV was isolated from SLEV-positive mosquito pools collected across multiple regions of California (**Figure 1A**). Contemporary isolates all belong to genotype III, contrasting with an historical California genotype V strain from 2003 (2003_V) (**Figure 1B**). The 2003_V strain was included as a representative from the period preceding the 11-year gap in detected SLEV activity that began in 2004. We evaluated strain-specific fitness in three cell types (African green monkey kidney [Vero], duck embryonic fibroblast [DEF], and *Cx. tarsalis* [CxTr]), two colonized *Cx.* vector species (*Cx. tarsalis* and *Cx. quinquefasciatus*), and 1 lineage (Collaborative Cross RIX 024X023) of immunocompetent adult mice (**Figure 1C**). All virus strains were minimally passaged in Vero cells prior to use (**Table 1**).

**Figure 1:**
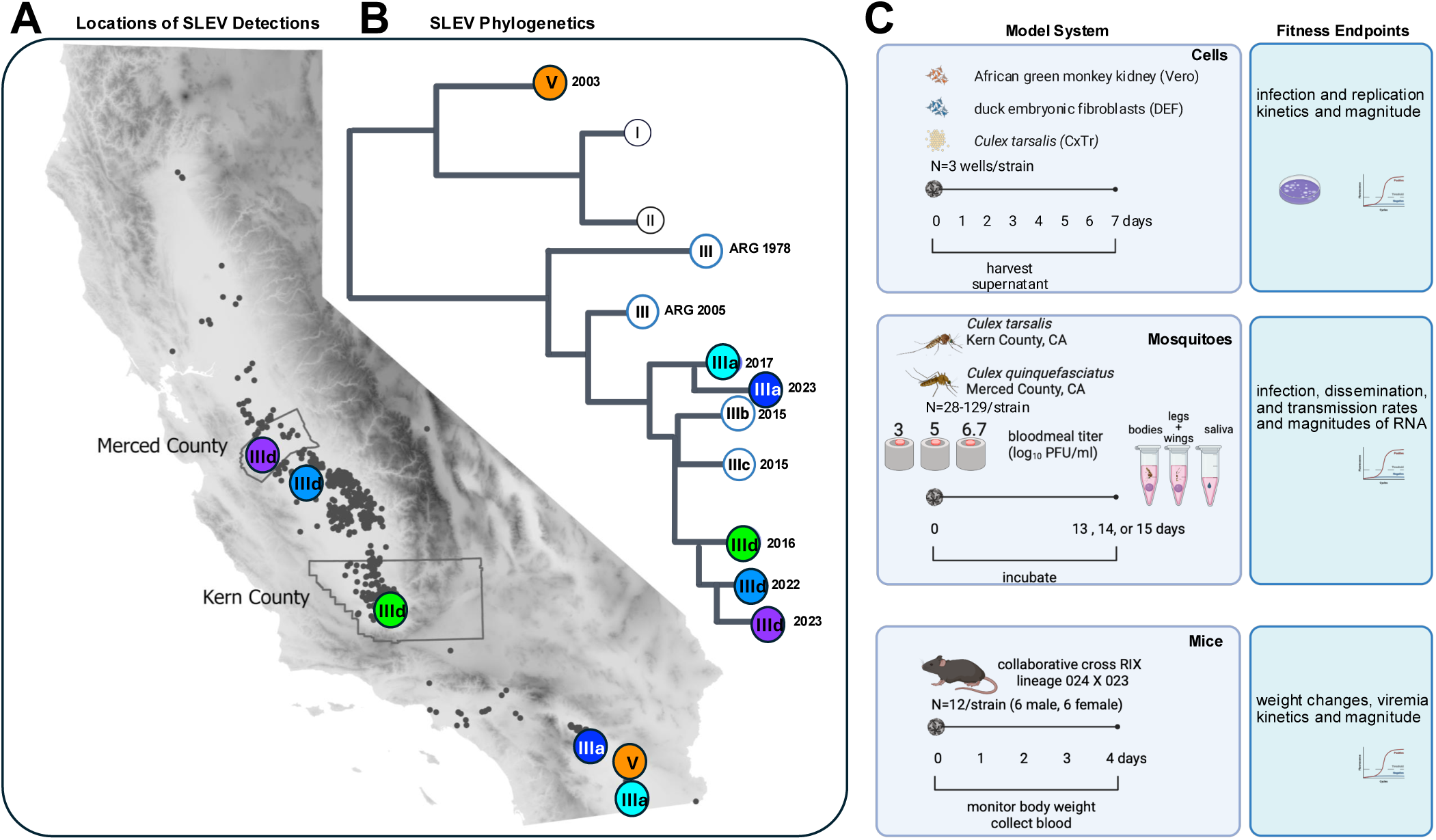
(**A**) Map of California showing locations of SLEV detections in mosquito pools from 2015-2025. The sources of the SLEV strains used in this study are shown in the colored circles and correspond to the labels on (**B**) the phylogenetic tree. ARG indicates Argentina. (**C**) Experimental design for SLEV studies in 3 model systems: cells, colonized *Cx. tarsalis* and *Cx. quinquefasciatus* mosquitoes from California, and collaborative cross RIX 024X023 mice. Fitness endpoints for each model are indicated. The image in C was created in BioRender (46).

### SLEV infection kinetics in cells

We first assessed fitness of SLEV in cells by measuring infection kinetics over 7 days (**Figure 2**). In Vero cells (**Figure 2A**), infection kinetics were similar among SLEV strains, with all viruses reaching mean peak titers of 7-8 log_10_ PFU/ml 3 dpi. Mean titers for most genotype III strains were comparable to 2003_V, although strain 2017_IIIa produced significantly higher titers at earlier time points, from 1-3 dpi [mixed effects 2-way ANOVA, p≤0.0001]. In DEF cells (**Figure 2B**), most strains also reached peak titers 3 dpi. All genotype III strains produced significantly higher mean DEF titers than 2003_V, particularly from 3-5 dpi [mixed effects 2-way ANOVA, p≤0.01], although no significant differences among strains were detected 6 dpi. From 2-4 dpi, all genotype III strains reached mean titers at least 1 log_10_ PFU/ml higher than 2003_V in DEF cells. In DEF cells, inter-strain variability in infection kinetics was greater than in Vero cells. In CxTr cells (**Figure 2C**), peak titers were delayed relative to the two vertebrate cell lines, with maximum titers 4 or 5 dpi. SLEV strains 2016_IIId and 2022_IIId produced significantly higher mean titers than 2003_V on multiple days [mixed effects 2-way ANOVA, p≤0.01], whereas 2017_IIIa produced lower titers compared to 2003_V at some times. Overall, these results indicate that some contemporary genotype III SLEV strains exhibit higher fitness than the historical 2003 genotype V strain in avian DEF and mosquito CxTr cells, while infection differences are less pronounced in mammalian Vero cells.

**Figure 2.**
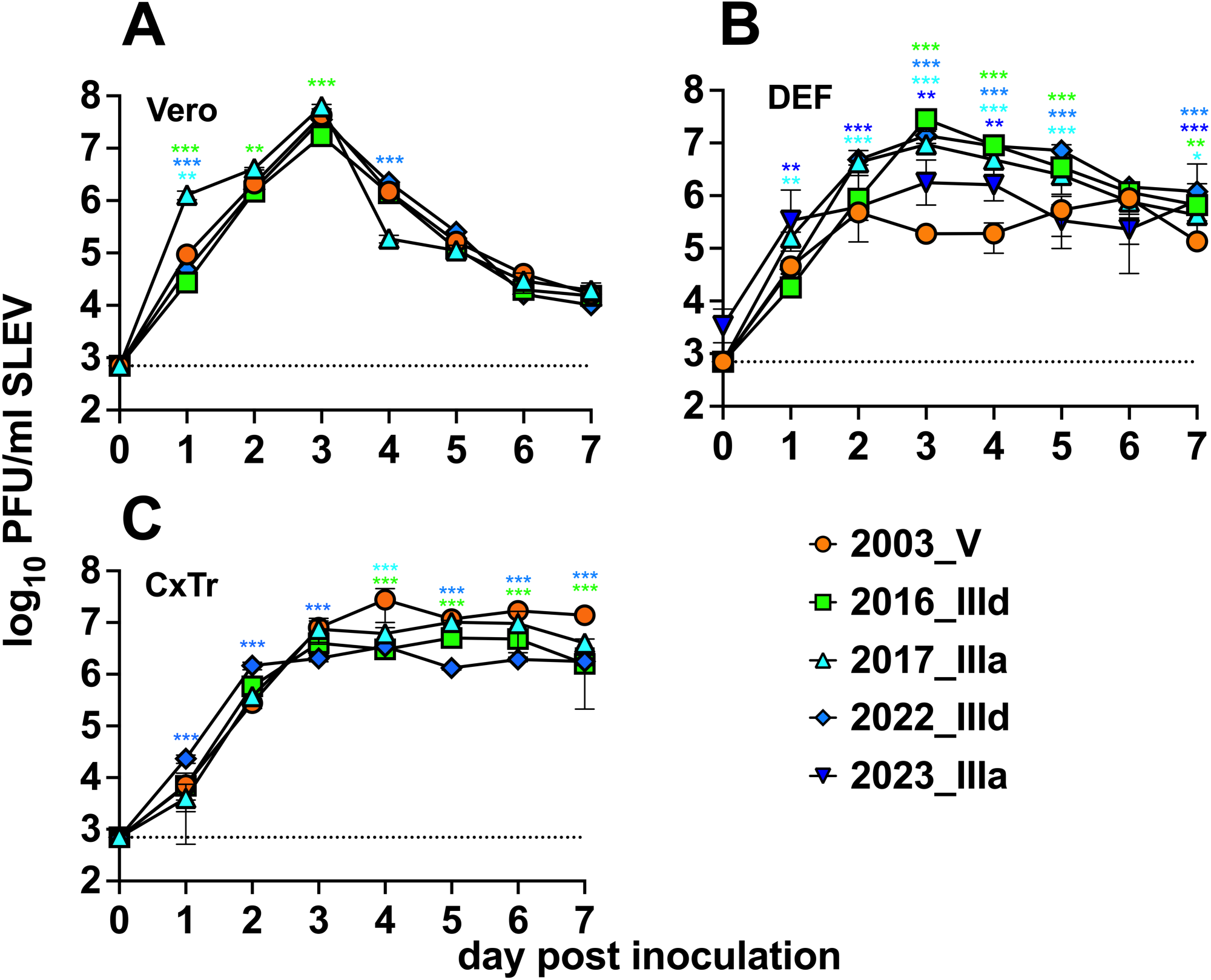
*In vitro* infection kinetics of SLEV strains in African green monkey kidney (Vero), duck embryonic fibroblast (DEF), and *Cx. tarsalis* (CxTr) cells. A multiplicity of infection (MOI) of 0.1 was used. The log transformed geometric mean virus titer from triplicate wells at each time point was plotted as PFU per milliliter. Error bars show geometric standard deviations. The dotted line represents the limit of detection, 2.8 log_10_ PFU/ml. Asterisks show comparisons of mean titers between 2003_V and other strains analyzed by mixed effects models with 2-way ANOVA on log-transformed values: *is p ≤0.01, **is p≤0.001, *** is p≤0.0001. Colors of asterisks show which strain is different compared to 2003_V.

### Dose response in mosquito vectors

We assessed the vector competence of the same five SLEV strains used in cell infections plus one additional 2023 strain (2023_IIId). We selected *Cx. tarsalis* and *Cx. quinquefasciatus* for evaluation because these species most frequently test positive for SLEV in California, representing 47% (1289/2731) and 48% (1316/2731) respectively, of SLEV mosquito pool detections statewide from 2015-2025 (**Supplemental Figure 2**). *Cx. tarsalis* were offered bloodmeals containing one of three SLEV doses, 3, 5, or 6.7 log_10_ PFU/ml, and *Cx. quinquefasciatus* were offered bloodmeals containing 5 or 6.7 log_10_ PFU/ml (**Figure 1C**). Infection (bodies), dissemination (legs & wings) and transmission (saliva) rates, calculated as the fraction of positive samples out of the total number of samples of that tissue tested multiplied by 100, increased with bloodmeal titer for both species (**Figure 3**, **Table 4**). At 3 log_10_ PFU/mL, infection rates were 0% or low for all strains in *Cx. tarsalis.* In contrast, infection, dissemination, and transmission rates increased when *Cx. tarsalis* ingested blood containing 5 or 6.7 log_10_ PFU/mL SLEV and in *Cx. quinquefasciatus* that ingested 6.7 versus 5 log_10_ PFU/ml. Back-titration confirmed that delivered bloodmeal doses were within ±0.5 log_10_ PFU/mL of the intended concentrations.

**Figure 3:**
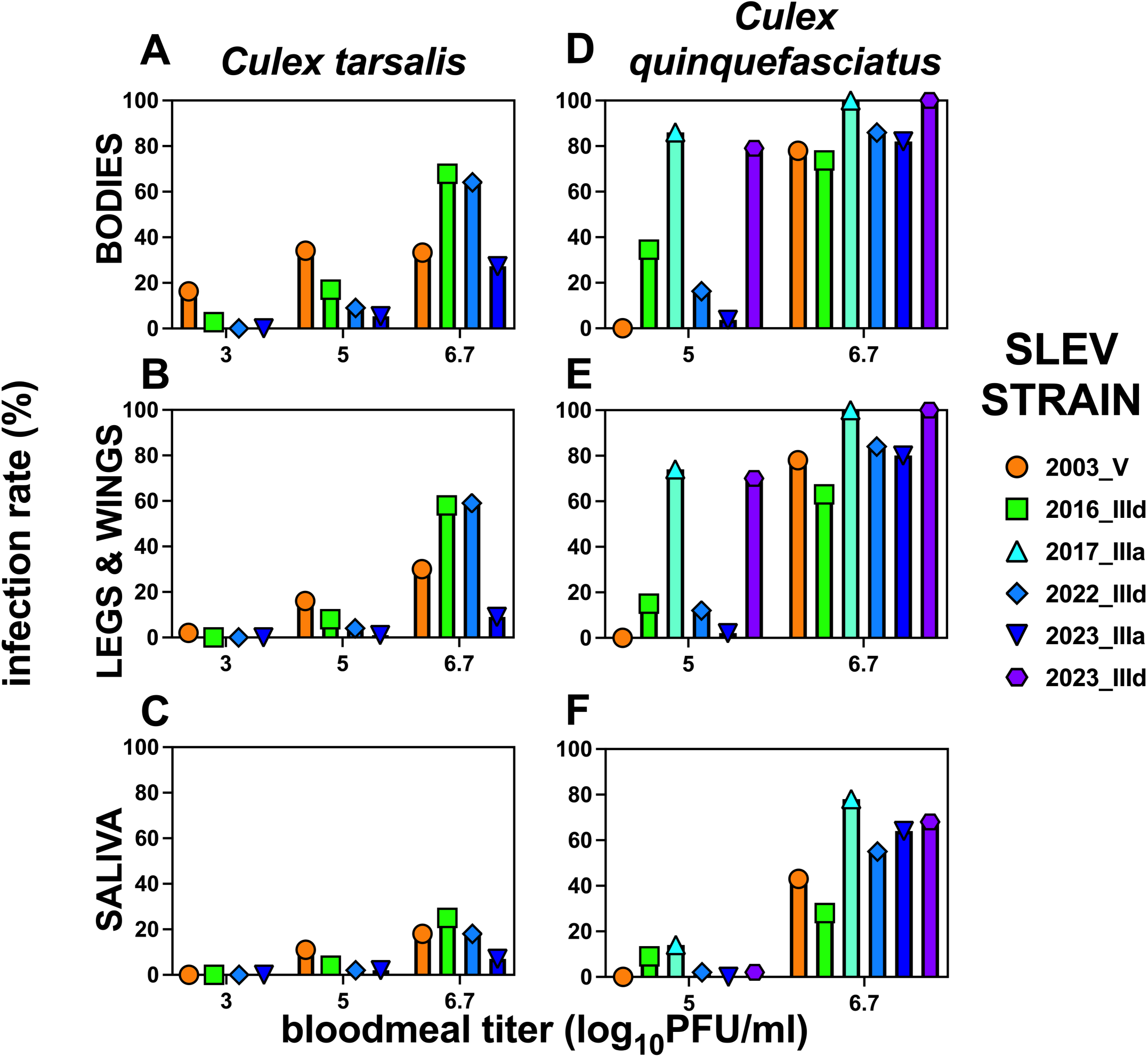
*Cx. tarsalis* and *Cx. quinquefasciatus* dose response to SLEV represented as infection, dissemination, and transmission rates 13, 14, or 15 days post-bloodmeal in cohorts of mosquitoes that ingested bloodmeals containing different SLEV strains. Infection rates were calculated as the number of positive bodies, legs & wings or saliva samples divided by the number of total samples of the same type that were tested, multiplied by 100. SLEV, St. Louis encephalitis virus; PFU, plaque-forming units.

**Table 4:**
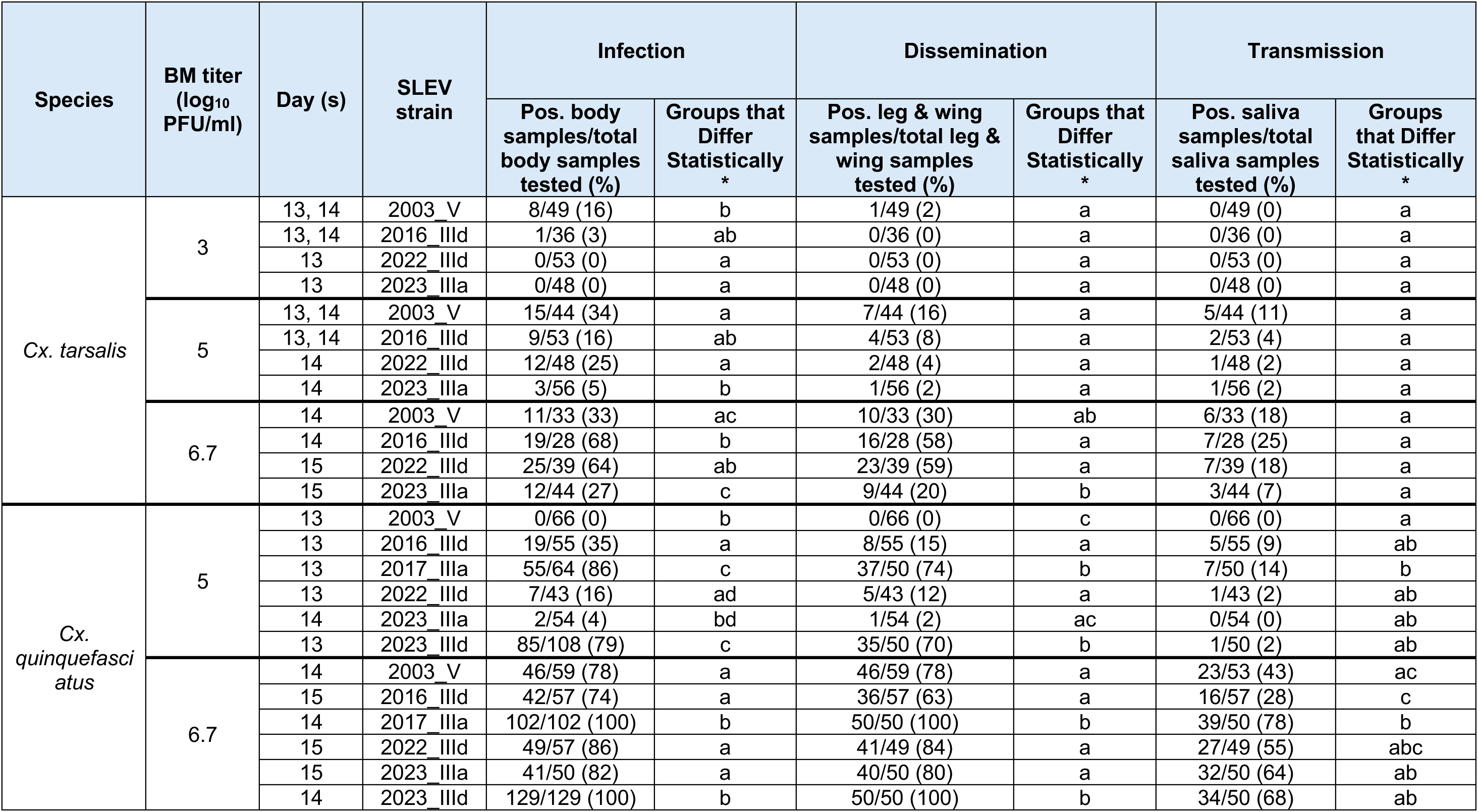
Fractions and rates (%) of mosquito body, leg & wing, and saliva samples that tested positive for SLEV RNA in colonized California *Cx. tarsalis* and *Cx. quinquefasciatus* mosquitoes that ingested bloodmeals containing different SLEV strains. *Letters show two-sided Fisher exact pairwise comparisons with Holm correction: values with no shared letter differ significantly at adjusted p<0.05; values sharing at least one letter are not significantly different. BM, bloodmeal, Pos., positive, SLEV, St. Louis encephalitis virus; PFU, plaque-forming units; *Cx*., *Culex*.

### SLEV strain comparisons in mosquito vectors

We compared the relative fitness of SLEV strains in both mosquito species (**Figure 3**, **Table 4**). In *Cx. tarsalis* at 3 log_10_ PFU/mL, all strains produced either no or low infection (**Figure 3A**), and dissemination (**Figure 3B**) and transmission (**Figure 3C**) were similarly rare. At 5 log_10_ PFU/mL, infection rates of the 2003 genotype V strain were significantly higher than those of 2023_IIIa while infection rates for 2003_V did not differ from those of 2016_IIId or 2022_IIId. The infection rate of 2022_IIId was also significantly higher than 2023_IIIa. Dissemination and transmission rates at this dose did not differ significantly among strains. At 6.7 log_10_ PFU/mL, infection rates of 2016_IIId were significantly higher than 2003_V and 2023_IIIa. No significant differences were observed in dissemination or transmission rates at this dose. Across doses, the two IIId lineage strains (2016_IIId and 2022_IIId), exhibited similar fitness profiles in all *Cx. tarsalis* tissues. Overall, these findings show strain-specific differences in SLEV fitness in *Cx. tarsalis*, with some but not all genotype III strains demonstrating higher infection rates than the historical genotype V strain, but without consistent differences in dissemination or transmission.

In *Cx. quinquefasciatus*, at 5 log_10_ PFU/ml no mosquitoes became infected after ingesting the 2003 genotype V strain. In contrast, all five genotype III strains produced infection (**Figure 3D**), ranging from 4% to 85%. At the 5 log_10_ PFU/ml dose, infection rates of 2017_IIIa were significantly higher than those of 2016_IIId and 2023_IIIa. Dissemination rates (**Figure 3E**) for all strains except 2023_IIIa were higher than 2005_V. Strain 2017_IIIa disseminated significantly better than 2016_IIId, 2022_IIId and 2023_IIIa. Only 2017_IIIa transmitted (**Figure 3F**) significantly better than 2003_V. At 6.7 log_10_ PFU/mL, most *Cx. quinquefasciatus* mosquitoes exposed to any SLEV strain became infected, and numerous statistically significant differences were observed. The 2017_IIIa and 2023_IIId strains, which infected 100% of bodies, produced significantly higher infection rates than 2003_V. At the 6.7 log_10_ PFU/ml dose, dissemination rates for all strains were also high (63–100%), and SLEV RNA in saliva was also frequently detected (28–78%). Dissemination rates were significantly higher for 2017_IIIa and 2023_IIId compared to all other strains. Overall, genotype III SLEV strains demonstrated high and strain-specific fitness in both *Cx.* vector species, with several strains infecting and transmitting more efficiently than the historical 2003 genotype V strain. Despite this, there was no trend of increasing fitness associated with the year a strain was detected.

### SLEV RNA levels in mosquito tissues

We quantified SLEV RNA levels in individual infected mosquitoes from both species (**Figure 4**). Mean titers decreased in the order bodies > legs & wings > saliva. In *Cx. tarsalis* at 5 log_10_ PFU/mL, mean RNA levels in bodies were significantly higher for 2016_IIId compared with 2022_IIId and 2023_IIIa (**Figure 4A**). No significant differences in mean RNA levels were detected among strains in legs and wings or saliva at this dose. At 6.7 log_10_ PFU/mL, mean RNA levels in *Cx. tarsalis* did not differ significantly among strains in any tissue type (**Figure 4B**). In *Cx. tarsalis*, mean RNA levels in any tissue for 2003_V were not significantly different compared to any other SLEV strain at both virus doses.

**Figure 4:**
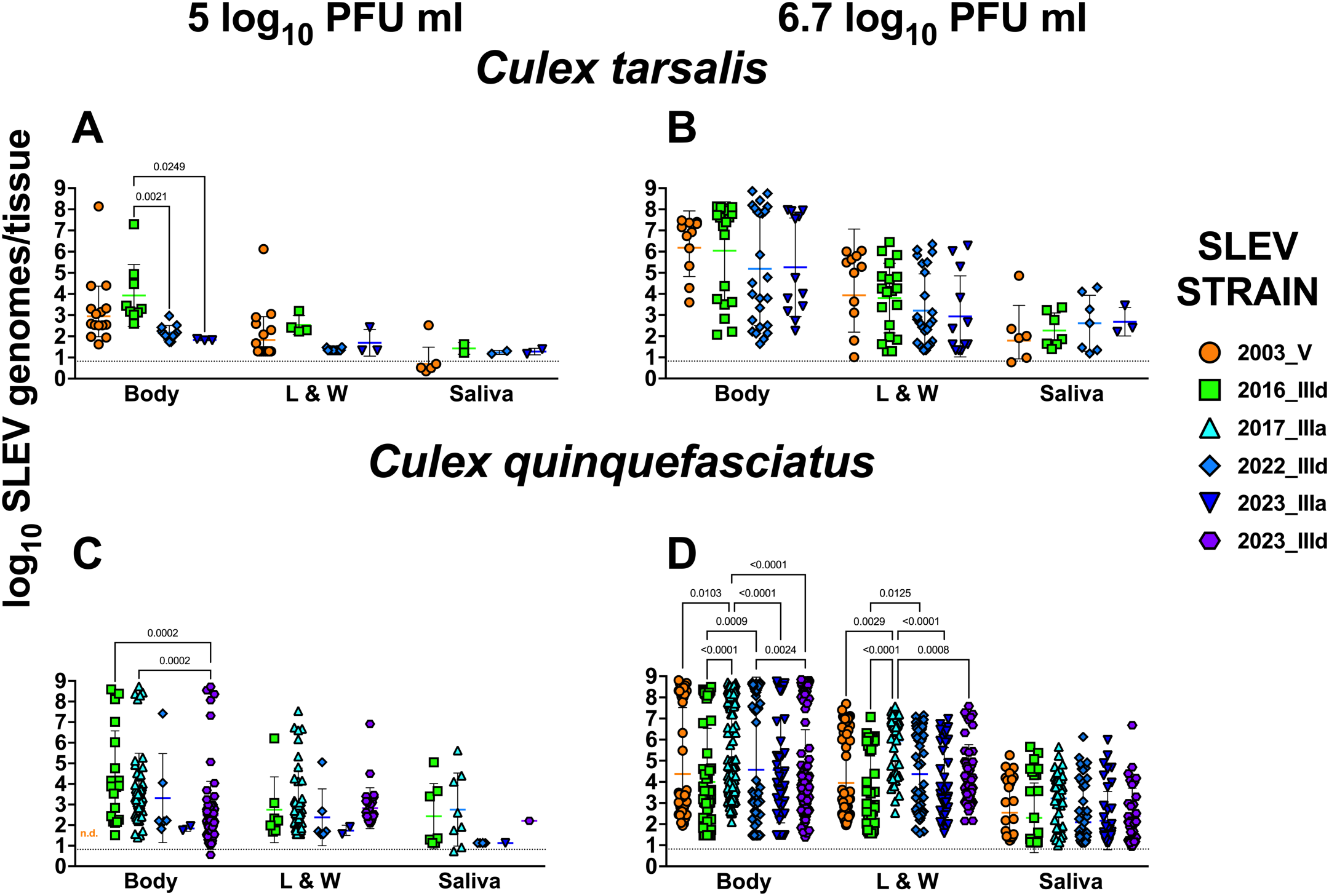
SLEV RNA levels 13,14, or 15 days post-bloodmeal in individual *Cx. tarsalis* and *Cx. quinquefasciatus* mosquitoes that ingested 5 or 6.7 log_10_ PFU/mL of different SLEV strains. L & W represents legs & wings. Levels were not statistically significantly different unless indicated with a p-value (Two-way ANOVA, Šídák’s multiple comparisons test; only values with p ≤ 0.05 were considered significant). Solid horizontal lines show geometric means and whiskers show geometric standard deviations. The dotted line shows the mean limit of detection (LOD), 0.8 log_10_ genomes/tissue. The LOD for some samples with a detectable measurement was below the mean LOD, which explains why some samples are under the LOD line.

In *Cx. quinquefasciatus* at 5 log_10_ PFU/mL, mean RNA levels in bodies were significantly higher for 2016_IIId and 2017_IIIa compared with 2023_IIId (**Figure 4C**). At 6.7 log_10_ PFU/mL, differences were observed across multiple strains in bodies and legs and wings (**Figure 4D**) where mean RNA levels for 2017_IIIa were significantly higher than all other strains. No differences in mean RNA levels were detected in saliva from any strain at either dose. In *Cx. quinquefasciatus* at 6.7 log_10_ PFU/mL, mean RNA levels in bodies and legs and wings for 2003_V were significantly lower than 2017_IIIa but were not statistically different from any other SLEV strain. Together, these data show that SLEV RNA detection was tissue- and strain-dependent, with the highest viral RNA levels detected in bodies, limited strain-specific differences in *Cx. tarsalis*, more pronounced differences in *Cx. quinquefasciatus* (particularly at the highest dose), and no significant strain-associated variation in saliva RNA levels.

### SLEV infection detection in saliva

We next attempted detection of infectious SLEV in mosquito saliva (**Table 6**) as a surrogate for transmission to a vertebrate host, since measurement of SLEV RNA alone does not indicate infectivity. Undiluted saliva samples from a randomly selected subset of SLEV RNA positive *Cx. quinquefasciatus* that ingested 6.7 log_10_ PFU/mL of either 2017_IIIa or 2023_IIId were tested using qualitative titrations. Plaque formation was observed in all saliva samples with SLEV RNA levels above 1.5 log_10_ genomes per sample. These results indicate that *Cx. quinquefasciatus* is capable of transmitting infectious genotype III SLEV.

### Vector species comparisons

Using the same data from Figure 3 and Table 4, we evaluated the relative fitness of SLEV across the two *Cx*. species by comparing infection, dissemination, and transmission rates at matched bloodmeal doses for the four strains that were evaluated in both species (**Figure 5**). At 5 log_10_ PFU/ml, *Cx. quinquefasciatus* showed no susceptibility (0%, 0/66) to the 2003 genotype V strain, whereas *Cx. tarsalis* exhibited infection, dissemination, and transmission rates of 34% (15/44), 15% (7/44), and 11% (5/44), respectively. In contrast, for the three genotype III strains, infection, dissemination, and transmission rates at 5 log_10_ PFU/mL were not statistically different between the two species. At 6.7 log_10_ PFU/mL, infection rates in *Cx. quinquefasciatus* trended (2016_IIId, 2022_IIId) or were significantly (2003_V, 2023_IIIa) higher than in *Cx. tarsalis*. Dissemination and transmission rates also followed this same pattern where rates were higher in *Cx. quinquefasciatus* compared to *Cx. tarsalis*. Overall, these results indicate that the SLEV strains tested exhibit comparable or greater fitness in *Cx. quinquefasciatus* than *Cx. tarsalis*.

**Figure 5:**
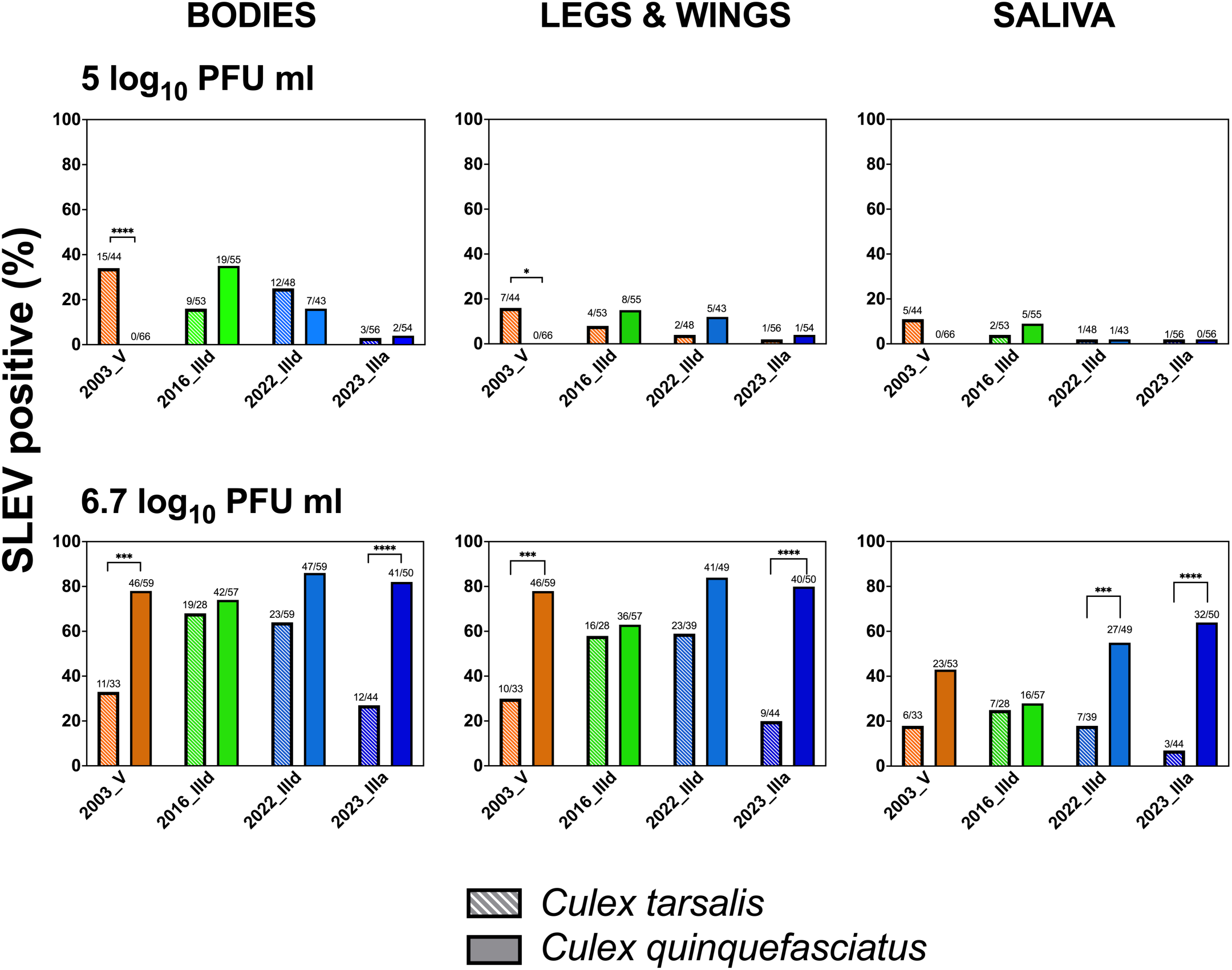
Rates of SLEV infection (bodies), dissemination (legs & wings), and transmission (saliva) by species 13, 14, or 15 days post-bloodmeal in cohorts of *Cx. tarsalis* and *Cx. quinquefasciatus* mosquitoes that ingested 5 or 6.7 log_10_ PFU/mL of different SLEV strains. Rates were calculated as the number of positive bodies, legs & wings or saliva samples divided by the number of total samples of the same type that were tested, multiplied by 100. Rates were not statistically significantly different unless indicated with a symbol (Fisher’s exact tests; only values p ≤ 0.05 were considered significant, *** is p<0.001, **** is p<0.00001). SLEV, St. Louis encephalitis virus; PFU, plaque-forming units; *Cx*., *Culex*.

### SLEV genetic determinants of vector fitness

We compared genome-wide amino acid differences across strains (**Table 5**) and via alignment (**Supplemental Figure 3**) to evaluate whether variant amino acids are shared between strains with similar vector fitness (e.g. 2017_IIIa and 2023_IIId that infected the majority of *Cx. quinquefasciatus* at both 5 and 6.7 log_10_ PFU/ml doses). Of 60 amino acid differences across any of the six strains, 38 were unique to the 2003 V strain. There were no shared amino acid differences in strains with high (2017_IIIa and 2023_IIId) or low (2016_IIId, 2022_IIId, 2023_IIIa) *Cx. quinquefasciatus* infection patterns.

**Table 5:** Amino acid differences (N=60) across 6 SLEV strains used for this study. Grey shading shows the minority amino acid(s). Charged residues are indicated in colors where **K/R/H are blue** and **D/E are red**. X denotes incomplete sequence in GenBank record. ins indicates insertion. C is capsid, prM is premembrane, M is membrane, E is envelope, NS is non-structural.

| Gene | Genomic amino acid Position | Gene amino acid position | 2003_V | 2016_IIId | 2017_IIIa | 2022_IIId | 2023_IIIa | 2023_IIId |
| --- | --- | --- | --- | --- | --- | --- | --- | --- |
| C | 10 | 10 | R | K | K | K | K | K |
|  | 71 | 71 | A | A | V | A | A | A |
|  | 92 | 92 | L | M | L | L | L | L |
|  | 114 | 114 | A | A | A | V | A | A |
|  | 116 | 116 | I | I | I | T | I | I |
| prM | 261 | 142 | A | V | V | V | V | V |
| E | 340 | 52 | D | E | E | E | E | E |
|  | 352 | 64 | T | S | S | S | S | S |
|  | 450 | 162 | N | N | N | N | S | N |
| NS1 | 796 | 7 | D | N | D | D | D | D |
|  | 859 | 70 | N | S | S | S | S | S |
|  | 862 | 73 | R | K | K | K | K | K |
|  | 885 | 96 | H | Y | Y | Y | Y | Y |
|  | 887 | 98 | R | K | K | K | K | K |
|  | 891 | 102 | Q | R | R | R | R | R |
|  | 901 | 112 | D | N | N | N | N | N |
|  | 963 | 174 | G | E | E | E | E | E |
|  | 1077 | 288 | T | T | T | T | I | T |
| NS2A | 1236 | 33 | L | F | F | F | F | F |
|  | 1250 | 47 | I | V | I | I | I | I |
|  | 1291 | 88 | C | Y | Y | Y | Y | Y |
|  | 1295 | 92 | I | I | I | I | F | I |
|  | 1347 | 144 | A | A | A | A | V | A |
| NS2B | 1416 | 48 | K | R | R | R | R | R |
| NS3 | 1837 | 338 | A | A | A | T | A | A |
|  | 1853 | 354 | S | N | N | N | N | N |
|  | 1898 | 399 | K | K | K | K | R | K |
|  | 1978 | 479 | N | N | S | N | N | N |
|  | 2049 | 550 | V | V | V | A | V | A |
|  | 2088 | 589 | R | K | K | K | K | K |
|  | 2093 | 594 | R | K | K | K | K | K |
| NS4B | 2184 | 67 | T | T | T | T | T | A |
|  | 2195 | 78 | G | S | S | S | S | S |
|  | 2210 | 93 | L | S | S | S | S | S |
| NS5 | 2288 | 22 | G | G | E | G | E | G |
|  | 2289 | 23 | S | P | P | P | P | P |
|  | 2292 | 26 | F | F | F | F | F | S |
|  | 2293 | 27 | A | - | - | - | - | - |
|  | 2298 | 32 | P | S | P | S | P | S |
|  | 2299_ins | 33_ins | - | X | X | - | - | - |
|  | 2300 | 34 | D | X | X | D | D | D |
|  | 2332 | 66 | Q | Q | Q | Q | Q | H |
|  | 2343 | 77 | A | S | S | S | S | S |
|  | 2395 | 129 | V | I | I | I | I | I |
|  | 2418 | 152 | A | V | A | A | A | A |
|  | 2513 | 247 | V | I | I | I | I | I |
|  | 2649 | 383 | Y | H | H | H | H | H |
|  | 2692 | 426 | T | A | A | A | A | A |
|  | 2701 | 435 | T | I | I | I | I | I |
|  | 2705 | 439 | I | V | V | V | V | V |
|  | 2712 | 446 | M | T | T | T | T | T |
|  | 2939 | 673 | L | F | F | F | F | F |
|  | 2966 | 700 | E | K | K | K | K | K |
|  | 2988 | 722 | M | T | T | T | T | T |
|  | 3025 | 759 | N | S | S | S | S | S |
|  | 3030 | 764 | H | Y | Y | Y | Y | Y |
|  | 3119 | 853 | V | I | I | I | I | I |
|  | 3161 | 895 | V | I | I | I | I | I |
|  | 3199 | 933 | I | T | T | T | T | T |
|  | 3278 | 1012 | K | R | R | R | R | R |
|  | 3413 | 1147 | I | V | V | V | V | A |
|  | 3429 | 1163 | V | A | A | A | A | A |

**Table 6:**
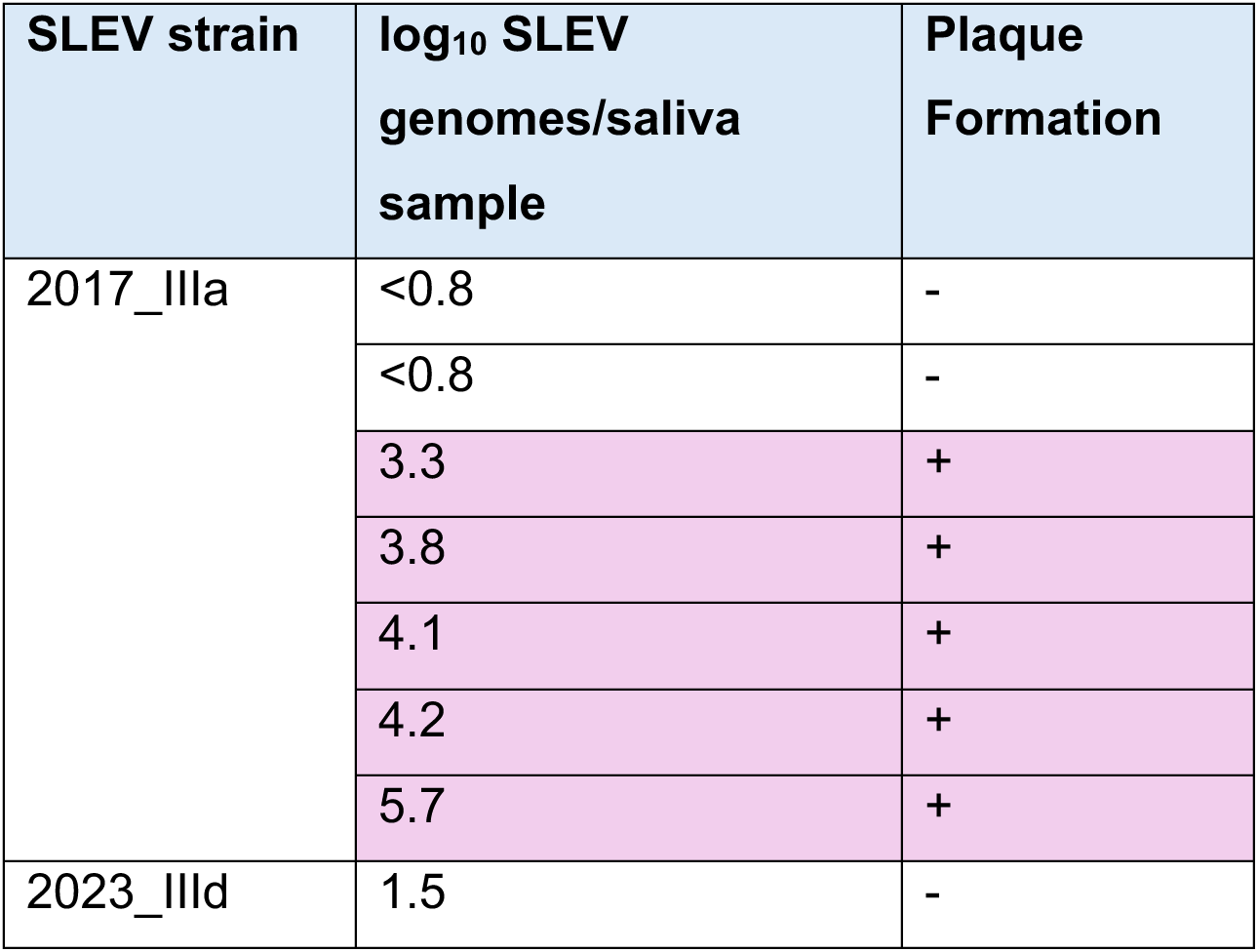
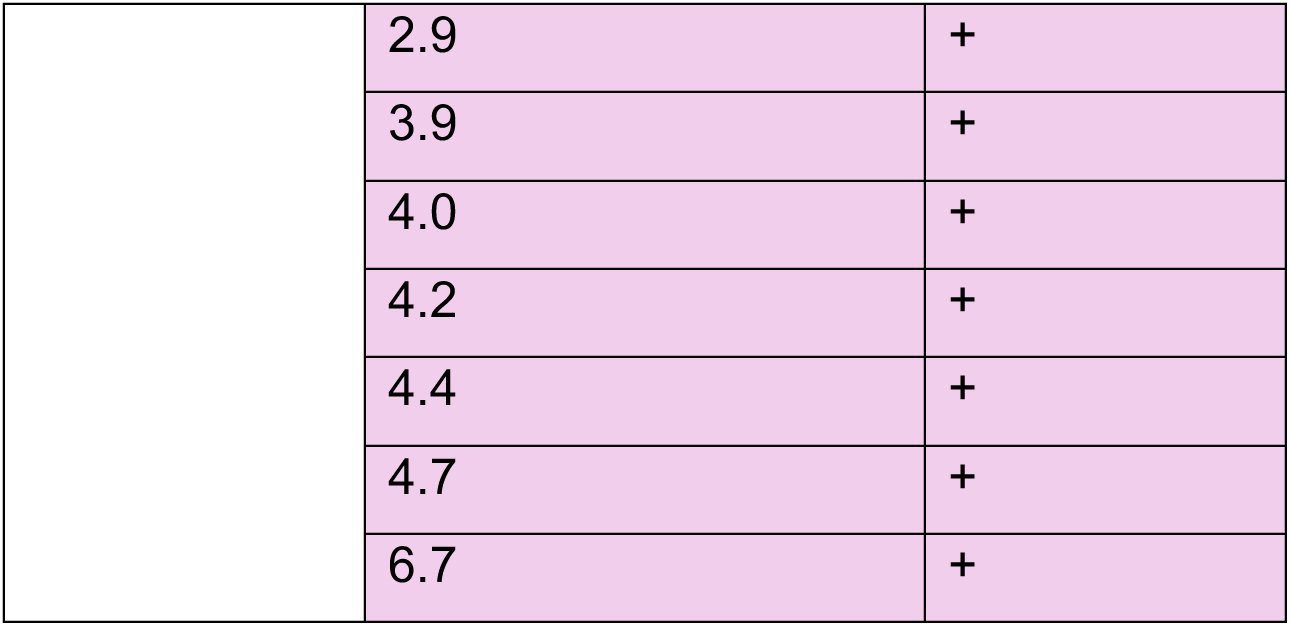
Infectious SLEV in *Cx. quinquefasciatus* saliva from mosquitoes that ingested 6.7 log_10_ PFU/mL in bloodmeals.

### SLEV kinetics in mice

We next evaluated SLEV viremia kinetics in mice. SLEV maintenance in nature requires transmission from viremic vertebrate hosts to mosquitoes. Immunocompetent collaborative cross RIX mice serve as a model of flavivirus infection. The RIX 024×023 lineage was selected for this study because WNV-inoculated mice of this lineage consistently develop detectable viremia (33, 34). None of the SLEV strains caused clinical disease in RIX 024X023 mice, and body weights remained within 5% of starting weight through the study endpoint at 4 dpi, similar to the mock-inoculated group (**Figure 6A**). All six SLEV strains produced detectable SLEV RNA in blood from 1-4 dpi, with peak RNA levels 2 or 3 dpi (**Figure 6B**). The historical 2003_V strain peaked at 2 dpi, whereas most genotype III strains peaked at 3 dpi. Despite this difference in peak timing, mean SLEV RNA levels did not vary significantly between any strain and 2003_V at 1, 2, or 4 dpi. At 3 dpi, however, mean RNA levels for 2022_IIId and 2023_IIId were significantly higher than those for 2003_V. Some differences were also observed among genotype III strains: at 1 dpi 2016_IIId and 2017_IIIa were significantly higher than 2023_IIId, whereas at 4 dpi 2023_IIId had significantly higher RNA levels than 2017_IIIa, 2022_IIId, and 2023_IIIa. Overall, these data indicate that all SLEV strains produce subclinical infection in collaborative cross RIX 024X023 mice, with acute viremia and similar RNA viremia kinetics, although most genotype III strains show a 1-day delay to peak viremia compared to the genotype V strain.

**Figure 6.**
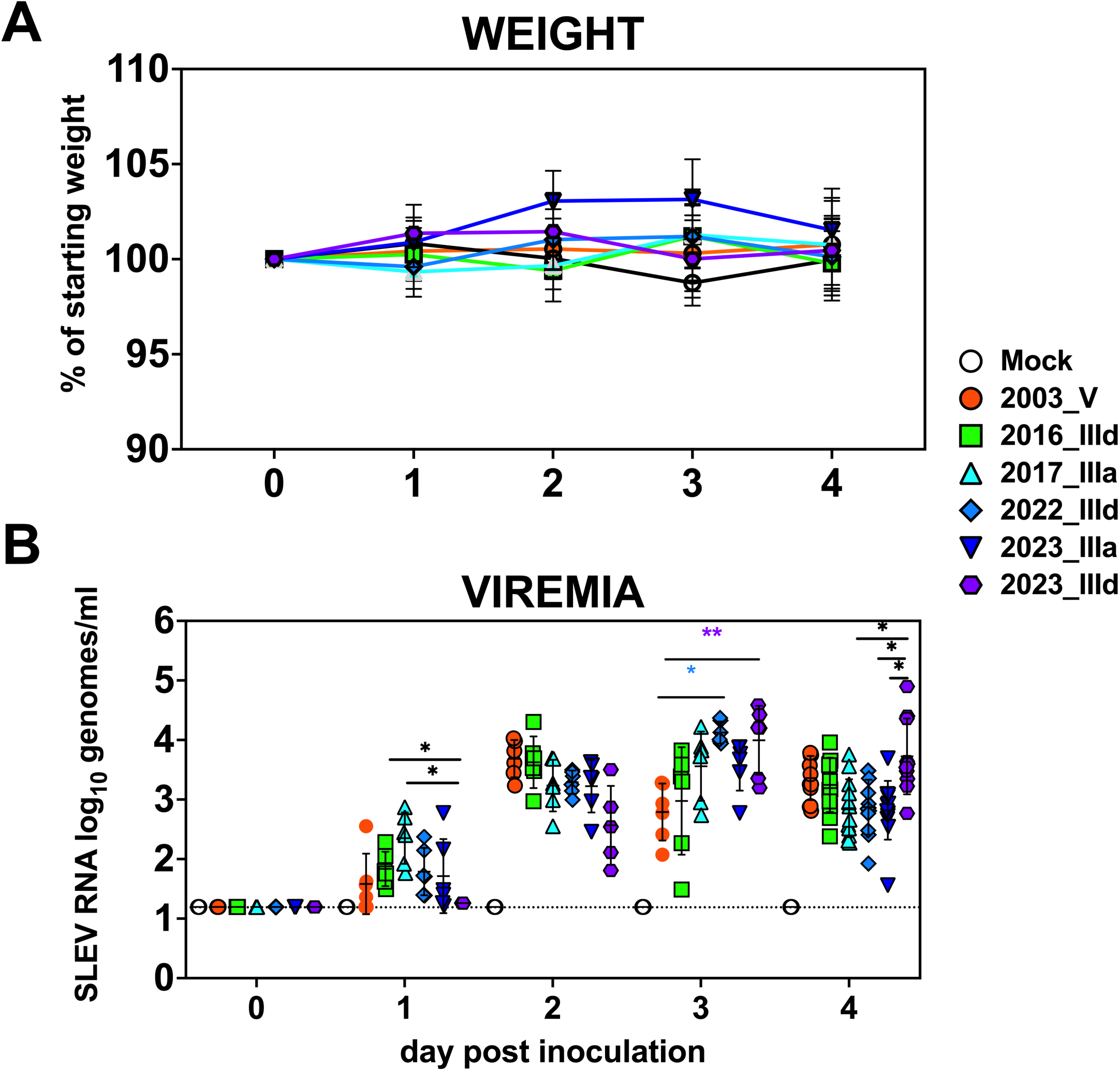
Comparison of SLEV strains in collaborative cross recombinant intercross (RIX) 024X023 mice. Shown are (**A**) percent weight change and (**B**) viremia from 0-4 days post-inoculation. Each group included six male and six female mice. For panel B, viremia is presented as log-transformed geometric mean RNA levels in blood where each symbol shows an individual animal. To minimize blood collection from individuals, 6 mice were sampled on days 1 and 3, while a different set of 6 mice was sampled on day 2. All 12 mice were sampled on day 4, immediately before euthanasia. Error bars indicate geometric standard deviations. The dotted horizontal line denotes the limit of detection, 1.2 log_10_ SLEV genomes/ml blood. Statistical comparisons between strains were performed using mixed-effects models with two-way ANOVA on log-transformed values: *is p ≤0.01, **is p≤0.001. n.d. is not detected. Data shown are from one experiment. Colored asterisks show which strain is different compared to 2003_V. Black asterisks show comparisons between genotype III strains.

## DISCUSSION

SLEV reemergence in California after 2015 represents a striking departure from historical patterns, in which activity was sporadic, spatially limited, and often disappeared for years following introduction. The persistence and geographic expansion of genotype III SLEV across many California counties since its reemergence in 2015 raises questions about whether increased SLEV fitness may be contributing to this altered geographic pattern. By comparing contemporary California SLEV strains from 2016-2023 with a historical genotype V strain from 2003 that circulated prior to WNV invasion and the 11-year disappearance of SLEV statewide from 2004-2014, our study provides the first experimental evidence evaluating fitness of post-2015 SLEV in cells, mosquito vectors, and mice.

Several of the five post-2015 genotype III strains we evaluated showed equal or greater infectivity than the historical 2003 genotype V strain in duck embryonic fibroblast and *Cx. tarsalis* cells and in *Cx. quinquefasciatus*, particularly at the bloodmeal dose of 5 log_10_ PFU/mL. The 2003 strain and most genotype III strains achieved similar magnitudes of SLEV RNA in mosquito tissues. Notably, *Cx. quinquefasciatus* exposed to the 2003 strain at the 5 log_10_ PFU/mL dose did not become infected; by contrast, each genotype III strain produced measurable infection and, for some strains, dissemination and SLEV RNA in saliva, our proxy for transmission. This differential suggests that some contemporary SLEV lineages may be better suited to infect *Cx. quinquefasciatus* than 2003_V under ecologically relevant viremia levels typical of passerine avian reservoirs, which range from 2-6 log_10_ PFU/ml (43). Detection of infectious SLEV in saliva from *Cx. quinquefasciatus* for the 2 evaluated genotype III SLEV strains also shows that *Cx. quinquefasciatus* can transmit infectious genotype III SLEV. Given that *Cx. quinquefasciatus* is abundant in urban areas of southern and central California (47, 48) which continue to report much of the SLEV activity statewide in mosquitoes (10), enhanced vector competence in this species may facilitate sustained and geographically widespread transmission. In contrast, *Cx. tarsalis* displayed more heterogeneous and strain-specific responses to SLEV. Some genotype III strains (notably 2016_IIId and 2022_IIId) showed higher infection, dissemination, and transmission success than the 2003 strain at the 6.7 log_10_ PFU/mL dose, although no clear hierarchy of fitness emerged among strains or related to genetic classification within genotype III or detection year. This complexity mirrors earlier work on WNV in California, where viral fitness differences in *Cx. tarsalis* were sometimes inconsistent across strains and relatively small compared to those observed in avian hosts (31, 32). Taken together, our results suggest that while *Cx. tarsalis* remains a competent vector for both historical and contemporary SLEV, the enhanced fitness of genotype III strains is more evident, and potentially more epidemiologically consequential, in *Cx. quinquefasciatus*.

In contrast with the data from mosquito vectors, we observed no major differences in SLEV viremia magnitudes in collaborative cross RIX 024×023 mice. Given that viremia is an important determinant of the likelihood of transmission to feeding mosquitoes, these results suggest that the strains evaluated here produce similar levels of vertebrate infectiousness. This finding suggests that phenotypic differences among strains may be more pronounced in mosquito vectors than in a vertebrate host. Future studies comparing SLEV strain fitness in avian reservoirs will be important for determining whether strain-specific differences in vertebrate amplification contribute to transmission dynamics in natural avian-mosquito cycling.

Another important outcome is the absence of a temporal fitness trend among post-2015 SLEV strains. Despite concerns that ongoing circulation of SLEV since 2015 might select for increasingly infectious genotypes, our data do not support a directional shift in viral fitness. Instead, fitness differences among genotype III strains appear idiosyncratic and not associated with year of detection. This pattern indicates that genetic variation within genotype III SLEV may influence fitness at a fine scale, but no single lineage among the strains we evaluated exhibits uniformly elevated infection or transmission potential. Furthermore, the lack of shared mutations in SLEV strains with higher vector fitness suggests that varying vector competence is likely driven by factors other than viral genetic differences in the SLEV strains evaluated, including shared combinations of amino acid changes, synonymous mutations, non-coding or RNA folding patterns, or intrahost virus population dynamics.

This study has limitations. Cell culture studies can identify differences in flavivirus replication, but they rely on single or few (for CxTr) cell types that may not be physiologically relevant for arbovirus infection. Monocultures also have altered or antiviral responses and do not capture the barriers to infection, dissemination, transmission, and immune evasion encountered in vectors and vertebrate hosts. Consequently, *in vitro* replication phenotypes may not accurately predict epidemiologically relevant fitness or transmission potential. We also used long-established laboratory colonies of *Cx. tarsalis* and *Cx. quinquefasciatus* that may not reflect the genetic or ecological variation of wild California mosquitoes. However, a prior study demonstrated that vector competence for WNV can vary over time within the same mosquito species (14), highlighting the value of using colonized mosquitoes derived from a single source population for direct comparisons of transmission phenotypes among virus strains. Local adaptation or hybridization within the *Cx. pipiens* complex could influence transmission outcomes for *Cx. quinquefasciatus* (which is part of the complex). Although collaborative cross RIX mice are more physiologically relevant than cell culture, they are not SLEV amplification hosts. Thus, viremia phenotypes may not accurately predict mosquito or maintenance of avian-mosquito transmission cycles, and because mice were euthanized 4 dpi, later viremia differences were not assessed. Although the SLEV strains evaluated represent the predominant genotype III lineages detected since 2015, limited sequencing of California SLEV since 2018 restricts our ability to assess whether these strains capture current viral diversity. Additional contemporary strains may display distinct vector fitness patterns. We also used a single genotype V comparator strain which does not represent all genotype V SLEV. Use of frozen rather than freshly cultured virus may have reduced infection (27), potentially underestimating infectivity, although comparisons among strains remain valid because all were tested under the same conditions. Mosquitoes exposed to different SLEV strains were harvested across a 1-day window (12-14 days post-exposure), but this is unlikely to have substantially affected infection or SLEV RNA levels based on relatively small kinetic changes in WNV over similar time periods (49–51). Saliva collected in capillary tubes provides only a surrogate measure of transmissibility because it does not replicate natural mosquito probing behavior or host factors influencing infection establishment. Vector competence experiments were conducted under controlled temperature and humidity conditions that do not reflect natural environmental variation, where temperature (50) and temperature variation (52) can alter infection dynamics. Except for a subset of saliva samples tested by infectious titrations, we quantified SLEV RNA rather than infectious virus in mosquitoes, potentially overestimating transmission potential if RNA detection does not correspond to infectivity (50). Finally, although dose-response experiments improve ecological relevance, artificial bloodmeal titers may not capture the full range of avian viremias; our (53) and others’ (54) work with other arboviruses and SLEV (55) show that artificial bloodmeals may misrepresent true infectiousness of vertebrate reservoirs for mosquitoes.

Our findings have implications for understanding SLEV ecology in California. The enhanced ability of some genotype III strains to infect *Cx. quinquefasciatus* may help explain why SLEV circulation has been concentrated in regions where this species is locally abundant, including the Coachella and Central Valleys of California. Besides increased mosquito vector competence, several other factors could contribute to SLEV reemergence. Expansion or movement of reservoir hosts and mosquito vectors into new areas, combined with environmental changes such as warmer temperatures or altered water availability that favor mosquito production and shorten the extrinsic incubation period, could increase transmission potential. Reemergence may also be driven by shifts in reservoir host demography, such as an influx of immunologically naïve birds, or by increased vertebrate competence in avian reservoirs. In addition, reduced cross-protection in WNV-immune birds could allow SLEV to circulate more effectively in bird populations. Finally, SLEV activity could reflect stochastic reintroduction and establishment events without major changes in viral traits or ecology, particularly if introduction coincides with favorable vector and host conditions.

This study provides phenotypic context for the genetic diversification of genotype III SLEV in the western US. As multiple genotype III lineages continue to circulate, defining how genetic variation translates into differences in virus fitness is important for forecasting areas of SLEV transmission and human disease risk. Future work based on more SLEV genomic data will integrate analyses of specific viral mutations that affect vector fitness and avian competence. Mixed-virus strain competition infections in mosquito vectors and avian reservoirs to minimize inter-host variability are also planned to further elucidate virological determinants underlying SLEV persistence in California in the post-WNV era.

## AUTHOR CONTRIBUTIONS

Conceptualization: MAFR, HL, LLC

Data Curation: MAFR, HL, LLC

Formal Analysis: MAFR, LLC

Funding Acquisition: LLC

Investigation: MAFR, HL, RL, EI, LLC

Methodology: MAFR, HL, SA, LLC

Project Administration: LLC

Resources: LLC

Supervision: LLC

Validation: MAFR, LLC

Visualization: MAFR, CMB, LLC

Writing-Original Draft Preparation: MAFR,LLC

Writing-Review and Editing: MAFR, HL, ET, RL, EI, SA, CMB, LLC

## ACKNOWLEDGEMENTS

We thank the staff from Fresno Westside Mosquito Abatement District, Coachella Valley Mosquito and Vector Control District, Kern Mosquito and Vector Control District, and Merced County Mosquito Abatement District for providing mosquito samples. We thank the Davis Arbovirus Research and Training surveillance team including Anil Singapuri and Sunny Lee for providing SLEV RNA detection data and SLEV positive mosquito pools for use in this project. We thank Rachel Lynch and Ginger Shaw at the UNC SGCF for their assistance breeding and procuring the mice. Figure 1C was created using BioRender.

## REFERENCES

1. Monath TP, Tsai TF. 1987. St. Louis encephalitis: Lessons from the last decade. American Journal of Tropical Medicine and Hygiene 10.4269/ajtmh.1987.37.40s.

2. Monath TP. 1980. Epidemiology. St. Louis encephalitis. American Public Health Association, Washington D.C.

3. Diaz A, Coffey LL, Burkett-Cadena N, Day JF. 2018. Reemergence of St. Louis encephalitis virus in the Americas. Emerging Infectious Diseases 10.3201/eid2412.180372.

4. Reisen WK. 2003. Epidemiology of St. Louis encephalitis virus. Adv Virus Res 61:139–183.

5. Reisen W, Lothrop H, Chiles R, Madon M, Cossen C, Woods L, Husted S, Kramer V, Edman J. 2004. West Nile virus in California. Emerg Infect Dis 10:1369–1378.

6. Fang Y, Reisen WK. 2006. Previous infection with West Nile or St. Louis encephalitis viruses provides cross protection during reinfection in house finches. Am J Trop Med Hyg 75:480–485.

7. Reisen WK, Lothrop HD, Wheeler SS, Kennsington M, Gutierrez A, Fang Y, Garcia S, Lothrop B. 2008. Persistent West Nile virus transmission and the apparent displacement St. Louis encephalitis virus in southeastern California, 2003-2006. J Med Entomol 45:494–508.

8. Venkat H, Krow-Lucal E, Hennessey M, Jones J, Adams L, Fischer M, Sylvester T, Levy C, Smith K, Plante L, Komatsu K, Staples JE, Hills S. 2015. Concurrent Outbreaks of St. Louis Encephalitis Virus and West Nile Virus Disease - Arizona, 2015. MMWR Morb Mortal Wkly Rep 64:1349–1350.

9. VectorSurv- - Vectorborne Disease Surveillance System team. VectorSurv.

10. California Department of Public Health. Vector Borne Disease Section Annual Reports.

11. Goddard LB, Roth AE, Reisen WK, Scott TW. 2002. Vector competence of California mosquitoes for West Nile virus. Emerg Infect Dis 8:1385–1391.

12. Kilpatrick AM, Kramer LD, Jones MJ, Marra PP, Daszak P. 2006. West Nile virus epidemics in North America are driven by shifts in mosquito feeding behavior. PLoS Biol 4:e82.

13. Reisen WK, Fang Y, Martinez VM. 2005. Avian host and mosquito (Diptera: Culicidae) vector competence determine the efficiency of West Nile and St. Louis encephalitis virus transmission. Journal of Medical Entomology 10.1093/jmedent/42.3.367.

14. Reisen WK, Barker CM, Fang Y, Martinez VM. 2008. Does variation in Culex (Diptera: Culicidae) vector competence enable outbreaks of West Nile virus in California? Journal of Medical Entomology 10.1603/0022-2585(2008)45%5B1126:DVICDC%5D2.0.CO;2.

15. Reisen WK, Meyer RP, Milby MM, Presser SB, Emmons RW, Hardy JL, Reeves WC. 1992. Ecological observations on the 1989 outbreak of St. Louis encephalitis virus in the southern San Joaquin Valley of California. J Med Entomol 29:472–482.

16. Reisen WK, Wheeler SS. 2016. Surveys for Antibodies Against Mosquitoborne Encephalitis Viruses in California Birds, 1996-2013. Vector Borne Zoonotic Dis 16:264–282.

17. Reisen WK, Milby MM, Presser SB, Hardy JL. 1992. Ecology of mosquitoes and St. Louis encephalitis virus in the Los Angeles Basin of California, 1987-1990. J Med Entomol 29:582–598.

18. Diaz LA, Ré V, Almirón WR, Farías A, Vázquez A, Sanchez-Seco MP, Aguilar J, Spinsanti L, Konigheim B, Visintin A, García J, Morales MA, Tenorio A, Contigiani M. 2006. Genotype III Saint Louis encephalitis virus outbreak, Argentina, 2005. Emerging Infectious Diseases 12:1752–1754.

19. Beranek M, Torres C, Laurito M, Farías A, Contigiani M, Almirón W, Diaz A. 2024. Emergence of genotype III St. Louis encephalitis virus in the western United States potentially linked to a wetland in Argentina. Acta Trop 250:107088.

20. White GS, Symmes K, Sun P, Fang Y, Garcia S, Steiner C, Smith K, Reisen WK, Coffey LL. 2016. Reemergence of St. Louis Encephalitis Virus, California, 2015. Emerg Infect Dis 22:2185–2188.

21. Swetnam DM, Stuart JB, Young K, Maharaj PD, Fang Y, Garcia S, Barker CM, Smith K, Godsey MS, Savage HM, Barton V, Bolling BG, Duggal N, Brault AC, Coffey LL. 2020. Movement of St. Louis encephalitis virus in the western united states, 2014-2018. PLoS Neglected Tropical Diseases 10.1371/journal.pntd.0008343.

22. Ridenour CL, Cocking J, Poidmore S, Erickson D, Brock B, Valentine M, Roe CC, Young SJ, Henke JA, Hung KY, Wittie J, Stefanakos E, Sumner C, Ruedas M, Raman V, Seaton N, Bendik W, Hornstra O’Neill HM, Sheridan K, Centner H, Lemmer D, Fofanov V, Smith K, Will J, Townsend J, Foster JT, Keim PS, Engelthaler DM, Hepp CM. 2021. St. Louis Encephalitis Virus in the Southwestern United States: A Phylogeographic Case for a Multi-Variant Introduction Event. Front Genet 12:667895.

23. Kneubehl AR, Rehm DP, Curtis MW, Wimmer BM, Bolling B, Broussard A, Vela J, Rocha J, Templeton L, Vigilant M, Standlee C, Presley SM, Lopez JE, Ronca SE. 2026. Geographically Distinct Circulation of Genotype II and III St. Louis Encephalitis Virus, Texas, USA, 2009-2024. Emerg Infect Dis 32:521–532.

24. Maharaj PD, Bosco-Lauth AM, Langevin SA, Anishchenko M, Bowen RA, Reisen WK, Brault AC. 2018. West Nile and St. Louis encephalitis viral genetic determinants of avian host competence. PLoS Neglected Tropical Diseases 12:1–17.

25. Maharaj PD, Anishchenko M, Langevin SA, Fang Y, Reisen WK, Brault AC. 2012. Structural gene (prME) chimeras of St Louis encephalitis virus and West Nile virus exhibit altered in vitro cytopathic and growth phenotypes. J Gen Virol 93:39–49.

26. Patiris PJ, Oceguera 3rd LF, Peck GW, Chiles RE, Reisen WK, Hanson CV. 2008. Serologic diagnosis of West Nile and St. Louis encephalitis virus infections in domestic chickens. Am J Trop Med Hyg 78:434–441.

27. Maharaj PD, Bolling BG, Anishchenko M, Reisen WK, Brault AC. 2014. Genetic determinants of differential oral infection phenotypes of West Nile and St. Louis encephalitis viruses in Culex spp. mosquitoes. Am J Trop Med Hyg 91:1066–1072.

28. Reisen WK, Chiles RE, Martinez VM, Fang Y, Green EN. 2003. Experimental infection of California birds with western equine encephalomyelitis and St. Louis encephalitis viruses. J Med Entomol 40:968–982.

29. Reisen WK, Kramer LD, Chiles RE, Martinez VM, Eldridge BF. 2000. Response of house finches to infection with sympatric and allopatric strains of western equine encephalomyelitis and St. Louis encephalitis viruses from California. J Med Entomol 37:259–264.

30. Moudy RM, Meola MA, Morin LL, Ebel GD, Kramer LD. 2007. A newly emergent genotype of West Nile virus is transmitted earlier and more efficiently by *Culex* mosquitoes. Am J Trop Med Hyg 77:365–370.

31. Worwa G, Hutton AA, Brault AC, Reisen WK. 2019. Comparative fitness of West Nile virus isolated during California epidemics. PLoS Neglected Tropical Diseases 10.1371/journal.pntd.0007135.

32. Duggal NK, Bosco-Lauth A, Bowen RA, Wheeler SS, Reisen WK, Felix TA, Mann BR, Romo H, Swetnam DM, Barrett ADT, Brault AC. 2014. Evidence for Co-evolution of West Nile virus and House Sparrows in North America. PLoS Neglected Tropical Diseases 10.1371/journal.pntd.0003262.

33. Graham JB, Swarts JL, Wilkins C, Thomas S, Green R, Sekine A, Voss KM, Ireton RC, Mooney M, Choonoo G, Miller DR, Treuting PM, Pardo Manuel de Villena F, Ferris MT, McWeeney S, Gale M, Lund JM. 2016. A Mouse Model of Chronic West Nile Virus Disease. PLoS Pathog 12:e1005996.

34. Graham JB, Thomas S, Swarts J, McMillan AA, Ferris MT, Suthar MS, Treuting PM, Ireton R, Gale M, Lund JM. 2015. Genetic diversity in the collaborative cross model recapitulates human West Nile virus disease outcomes. mBio 6:e00493–00415.

35. Jasperse BA, Mattocks MD, Noll KE, Ferris MT, Heise MT, Lazear HM. 2023. Neuroinvasive Flavivirus Pathogenesis Is Restricted by Host Genetic Factors in Collaborative Cross Mice, Independently of Oas1b. J Virol 97:e0071523.

36. Rocha RF, Del Sarto JL, Gomes GF, Gonçalves MP, Rachid MA, Smetana JHC, Souza DG, Teixeira MM, Marques RE. 2021. Type I interferons are essential while type II interferon is dispensable for protection against St. Louis encephalitis virus infection in the mouse brain. Virulence 12:244–259.

37. Marques RE, Del Sarto JL, Rocha RPF, Gomes GF, Cramer A, Rachid MA, Souza DG, Nogueira ML, Teixeira MM. 2017. Development of a model of Saint Louis encephalitis infection and disease in mice. J Neuroinflammation 14:61.

38. California Department of Public Health. 2016. Weekly Reports.

39. California Department of Public Health. St. Louis encephalitis virus.

40. Brault AC, Fang Y, Reisen WK. 2015. Multiplex qRT-PCR for the Detection of Western equine encephalomyelitis, St. Louis encephalitis, and West Nile viral RNA in Mosquito Pools (Diptera: Culicidae). J Med Entomol 52:491–499.

41. Cornel AJ, Holeman J, Nieman CC, Lee Y, Smith C, Amorino M, Brisco KK, Barrera R, Lanzaro GC, Mulligan Iii FS. 2016. Surveillance, insecticide resistance and control of an invasive *Aedes aegypti* (Diptera: Culicidae) population in California. F1000Res 5:194.

42. McAbee RD, Kang K, Stanich MA, Christiansen JA, Wheelock CE, Inman AD, Hammock BD, Cornel AJ. 2004. Pyrethroid tolerance in *Culex pipiens pipiens* var *molestus* from Marin County, California. Pest Management Science 60:359–368.

43. Bosco-Lauth AM, Kooi K, Hawks SA, Duggal NK. 2025. Cross-protection between West Nile virus and emerging flaviviruses in wild birds. Am J Trop Med Hyg 112:657–662.

44. Suppiah J, Nadaraju S, Hamzah S, Chee HY. 2020. Viability of dengue virus in culture stocks is efficiently preserved by storage in diluted forms. Trop Biomed 37:282–287.

45. Madeira F, Madhusoodanan N, Lee J, Eusebi A, Niewielska A, Tivey ARN, Lopez R, Butcher S. 2024. The EMBL-EBI Job Dispatcher sequence analysis tools framework in 2024. Nucleic Acids Res 52:W521–W525.

46. Coffey L. 2026. BioRender.

47. Barker CM, Eldridge BF, Reisen WK. 2010. Seasonal abundance of *Culex tarsalis* and *Culex pipiens* complex mosquitoes (Diptera: Culicidae) in California. J Med Entomol 47:759–768.

48. Kothera L, Nelms B, Savage HM, Reisen WK. 2012. Complexity of the *Culex pipiens* complex in California. Proc Pap Annu Conf Mosq Vector Control Assoc Calif 80:1–3.

49. Eastwood G, Kramer LD, Goodman SJ, Cunningham AA. 2011. West Nile virus vector competency of *Culex quinquefasciatus* mosquitoes in the Galapagos Islands. Am J Trop Med Hyg 85:426–433.

50. Reisen WK, Fang Y, Martinez VM. 2006. Effects of temperature on the transmission of West Nile virus by *Culex tarsalis* (Diptera: Culicidae). J Med Entomol 43:309–317.

51. Anderson SL, Richards SL, Tabachnick WJ, Smartt CT. 2010. Effects of West Nile virus dose and extrinsic incubation temperature on temporal progression of vector competence in *Culex pipiens quinquefasciatus*. J Am Mosq Control Assoc 26:103–107.

52. McGregor BL, Kenney JL, Connelly CR. 2021. The effect of fluctuating incubation temperatures on West Nile virus infection in *Culex* mosquitoes. Viruses 13:1822.

53. Moore AJ, Van Rompay KKA, Louie W, Watanabe JK, An S, Leung R, Usachenko JL, Chu PN, Olstad KJ, McCoy CS, Campos RK, Weaver SC, Rossi SL, Coffey LL. 2025. Rhesus macaques model human Mayaro virus disease and transmit to Aedes aegypti mosquitoes 10.1101/2025.04.16.649064.

54. Hanley KA, Cecilia H, Azar SR, Moehn BA, Gass JT, Oliveira Da Silva NI, Yu W, Yun R, Althouse BM, Vasilakis N, Rossi SL. 2024. Trade-offs shaping transmission of sylvatic dengue and Zika viruses in monkey hosts. Nat Commun 15:2682.

55. Meyer RP, Hardy JL, Presser SB. 1983. Comparative vector competence of *Culex tarsalis* and *Culex quinquefasciatus* from the Coachella, Imperial, and San Joaquin Valleys of California for St. Louis encephalitis virus. Am J Trop Med Hyg 32:305–311.

