## Supplemental Figures 1-3 for "Comparative fitness of reemerging St. Louis encephalitis virus in vertebrate and mosquito cells, *Culex tarsalis* and *Culex quinquefasciatus* mosquitoes, and mice"

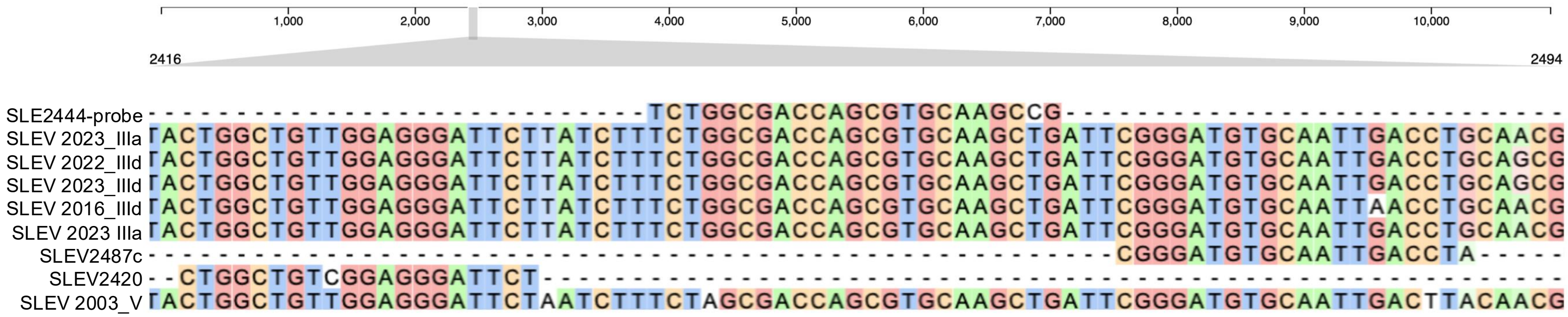


**Supplemental Figure 1**: SLEV multiple sequence alignment showing primer (SLEV2420, SLEV2487c) and probe (SLE2444-probe) binding regions including single nucleotide polymorphisms. SLEV strain names denote year of isolation and genotype.

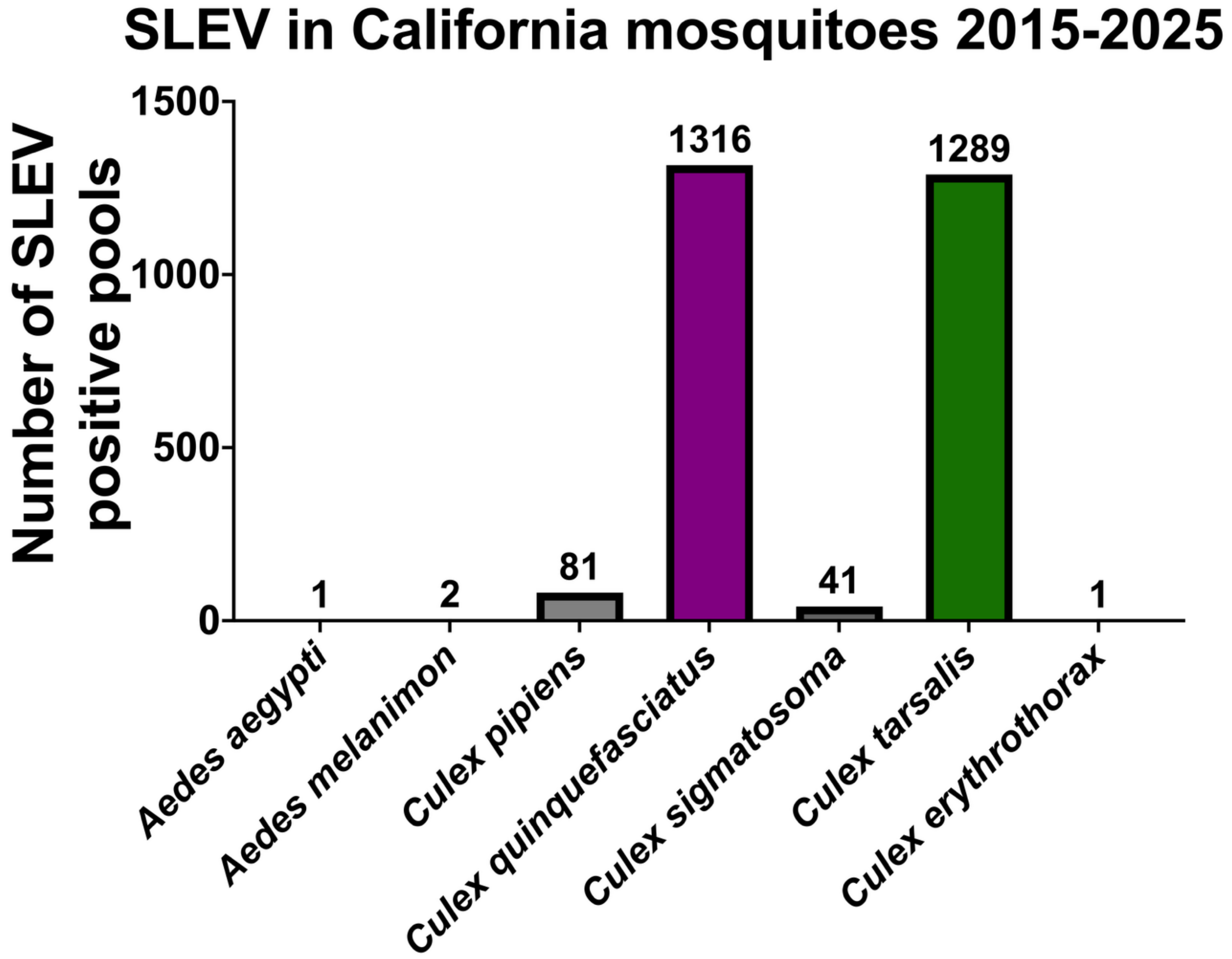


**Supplemental Figure 2**: SLEV positive mosquito pools in California from 2015-2025 by species. 2025 data is current as of December 8, 2025.

```
Capsid | prM
2003_V     MSKKPGKPGR NRVVNMLKRG VSRVNPLTGL KRILGSLLDG RGPVRFILAI LTFFRFTALQ PTEALKRRWR AVDKRTALKH LNGFKRDLGS MLDTINRRPS KKRGGTGSLL GLAALIGLAS SLQLSTYQGK VLMSINKTDA QSAINIPSAN  150
2016_IIId  MSKKPGKPGK NRVVNMLKRG VSRVNPLTGL KRILGSLLDG RGPVRFILAI LTFFRFTALQ PTEALKRRWR AVDKRTALKH LNGFKRDLGS MMDTINRRPS KKRGGTGSLL GLAALIGLAS SLQLSTYQGK VLMSINKTDA QSAINIPSAN  150
2017_IIIa  MSKKPGKPGK NRVVNMLKRG VSRVNPLTGL KRILGSLLDG RGPVRFILAI LTFFRFTALQ PTEALKRRWR VVDKRTALKH LNGFKRDLGS MLDTINRRPS KKRGGTGSLL GLAALIGLAS SLQLSTYQGK VLMSINKTDA QSAINIPSAN  150
2022_IIId  MSKKPGKPGK NRVVNMLKRG VSRVNPLTGL KRILGSLLDG RGPVRFILAI LTFFRFTALQ PTEALKRRWR AVDKRTALKH LNGFKRDLGS MLDTINRRPS KKRGGTGSLL GLAVLTGLAS SLQLSTYQGK VLMSINKTDA QSAINIPSAN  150
2023_IIIa  MSKKPGKPGK NRVVNMLKRG VSRVNPLTGL KRILGSLLDG RGPVRFILAI LTFFRFTALQ PTEALKRRWR AVDKRTALKH LNGFKRDLGS MLDTINRRPS KKRGGTGSLL GLAALIGLAS SLQLSTYQGK VLMSINKTDA QSAINIPSAN  150
2023_IIId  MSKKPGKPGK NRVVNMLKRG VSRVNPLTGL KRILGSLLDG RGPVRFILAI LTFFRFTALQ PTEALKRRWR AVDKRTALKH LNGFKRDLGS MLDTINRRPS KKRGGTGSLL GLAALIGLAS SLQLSTYQGK VLMSINKTDA QSAINIPSAN  150

prM | Envelope
2003_V     GANTCIVRAL DVGVMCKDDI TYLCPVLSAG NDPEDIDCWC DVEEVWVHYG RCTRMGHSRR SRRSISVQHH GDSTLATKNT PWLDTVKTTK YLTKVENWVL RNPGYALVAL AIGWMLGSNN TQRVVFVIML MLIAPAYSFN CLGTSNRDFV  300
2016_IIId  GANTCIVRAL DVGVMCKDDI TYLCPVLSAG NDPEDIDCWC DVEEVWVHYG RCTRMGHSRR SRRSISVQHH GDSTLATKNT PWLDTVKTTK YLTKVENWVL RNPGYALVAL VIGWMLGSNN TQRVVFVIML MLIAPAYSFN CLGTSNRDFV  300
2017_IIIa  GANTCIVRAL DVGVMCKDDI TYLCPVLSAG NDPEDIDCWC DVEEVWVHYG RCTRMGHSRR SRRSISVQHH GDSTLATKNT PWLDTVKTTK YLTKVENWVL RNPGYALVAL VIGWMLGSNN TQRVVFVIML MLIAPAYSFN CLGTSNRDFV  300
2022_IIId  GANTCIVRAL DVGVMCKDDI TYLCPVLSAG NDPEDIDCWC DVEEVWVHYG RCTRMGHSRR SRRSISVQHH GDSTLATKNT PWLDTVKTTK YLTKVENWVL RNPGYALVAL VIGWMLGSNN TQRVVFVIML MLIAPAYSFN CLGTSNRDFV  300
2023_IIIa  GANTCIVRAL DVGVMCKDDI TYLCPVLSAG NDPEDIDCWC DVEEVWVHYG RCTRMGHSRR SRRSISVQHH GDSTLATKNT PWLDTVKTTK YLTKVENWVL RNPGYALVAL VIGWMLGSNN TQRVVFVIML MLIAPAYSFN CLGTSNRDFV  300
2023_IIId  GANTCIVRAL DVGVMCKDDI TYLCPVLSAG NDPEDIDCWC DVEEVWVHYG RCTRMGHSRR SRRSISVQHH GDSTLATKNT PWLDTVKTTK YLTKVENWVL RNPGYALVAL VIGWMLGSNN TQRVVFVIML MLIAPAYSFN CLGTSNRDFV  300

Envelope
2003_V     EGASGATWID LVLEGGSCVT VMAPEKPTLD FKVMKMEATD LATVREYCYE ATLDTLSTVA RCPTTGEAHN TKRSDPTFVC KRDVVDRGWG NGCGLFGKGS IDTCAKFTCK NKATGKTILR ENIKYEVAIF VHGSTDSTSH GNYSEQIGKN  450
2016_IIId  EGASGATWID LVLEGGSCVT VMAPEKPTLD FKVMKMEATE LATVREYCYE ASLDTLSTVA RCPTTGEAHN TKRSDPTFVC KRDVVDRGWG NGCGLFGKGS IDTCAKFTCK NKATGKTILR ENIKYEVAIF VHGSTDSTSH GNYSEQIGKN  450
2017_IIIa  EGASGATWID LVLEGGSCVT VMAPEKPTLD FKVMKMEATE LATVREYCYE ASLDTLSTVA RCPTTGEAHN TKRSDPTFVC KRDVVDRGWG NGCGLFGKGS IDTCAKFTCK NKATGKTILR ENIKYEVAIF VHGSTDSTSH GNYSEQIGKN  450
2022_IIId  EGASGATWID LVLEGGSCVT VMAPEKPTLD FKVMKMEATE LATVREYCYE ASLDTLSTVA RCPTTGEAHN TKRSDPTFVC KRDVVDRGWG NGCGLFGKGS IDTCAKFTCK NKATGKTILR ENIKYEVAIF VHGSTDSTSH GNYSEQIGKN  450
2023_IIIa  EGASGATWID LVLEGGSCVT VMAPEKPTLD FKVMKMEATE LATVREYCYE ASLDTLSTVA RCPTTGEAHN TKRSDPTFVC KRDVVDRGWG NGCGLFGKGS IDTCAKFTCK NKATGKTILR ENIKYEVAIF VHGSTDSTSH GNYSEQIGKS  450
2023_IIId  EGASGATWID LVLEGGSCVT VMAPEKPTLD FKVMKMEATE LATVREYCYE ASLDTLSTVA RCPTTGEAHN TKRSDPTFVC KRDVVDRGWG NGCGLFGKGS IDTCAKFTCK NKATGKTILR ENIKYEVAIF VHGSTDSTSH GNYSEQIGKN  450

Envelope
2003_V     QAARFTISPQ APSFTANMGE YGTVTIDCEA RSGINTEDYY VFTVKEKSWL VNRDWFHDLN LPWTSPATTD WRNRETLVEF EEPHATKQTV VALGSQEGAL HTALAGAIPA TVSSSTLTLQ SGHLKCRAKL DKVKIKGTTY GMCDSAFTFS  600
2016_IIId  QAARFTISPQ APSFTANMGE YGTVTIDCEA RSGINTEDYY VFTVKEKSWL VNRDWFHDLN LPWTSPATTD WRNRETLVEF EEPHATKQTV VALGSQEGAL HTALAGAIPA TVSSSTLTLQ SGHLKCRAKL DKVKIKGTTY GMCDSAFTFS  600
2017_IIIa  QAARFTISPQ APSFTANMGE YGTVTIDCEA RSGINTEDYY VFTVKEKSWL VNRDWFHDLN LPWTSPATTD WRNRETLVEF EEPHATKQTV VALGSQEGAL HTALAGAIPA TVSSSTLTLQ SGHLKCRAKL DKVKIKGTTY GMCDSAFTFS  600
2022_IIId  QAARFTISPQ APSFTANMGE YGTVTIDCEA RSGINTEDYY VFTVKEKSWL VNRDWFHDLN LPWTSPATTD WRNRETLVEF EEPHATKQTV VALGSQEGAL HTALAGAIPA TVSSSTLTLQ SGHLKCRAKL DKVKIKGTTY GMCDSAFTFS  600
2023_IIIa  QAARFTISPQ APSFTANMGE YGTVTIDCEA RSGINTEDYY VFTVKEKSWL VNRDWFHDLN LPWTSPATTD WRNRETLVEF EEPHATKQTV VALGSQEGAL HTALAGAIPA TVSSSTLTLQ SGHLKCRAKL DKVKIKGTTY GMCDSAFTFS  600
2023_IIId  QAARFTISPQ APSFTANMGE YGTVTIDCEA RSGINTEDYY VFTVKEKSWL VNRDWFHDLN LPWTSPATTD WRNRETLVEF EEPHATKQTV VALGSQEGAL HTALAGAIPA TVSSSTLTLQ SGHLKCRAKL DKVKIKGTTY GMCDSAFTFS  600

Envelope
2003_V     KNPADTGHGT VIVELQYTGS NGPCRVPISV TANLMDLTPV GRLVTVNPFI STGGANNKVM IEVEPPFGDS YIVVGRGTTQ INYHWHKEGS SIGKALATTW KGAQRLAVLG DTAWDFGSIG GVFNSIGKAV HQVFGGAFRT LFGGMSWITQ  750
2016_IIId  KNPADTGHGT VIVELQYTGS NGPCRVPISV TANLMDLTPV GRLVTVNPFI STGGANNKVM IEVEPPFGDS YIVVGRGTTQ INYHWHKEGS SIGKALATTW KGAQRLAVLG DTAWDFGSIG GVFNSIGKAV HQVFGGAFRT LFGGMSWITQ  750
2017_IIIa  KNPADTGHGT VIVELQYTGS NGPCRVPISV TANLMDLTPV GRLVTVNPFI STGGANNKVM IEVEPPFGDS YIVVGRGTTQ INYHWHKEGS SIGKALATTW KGAQRLAVLG DTAWDFGSIG GVFNSIGKAV HQVFGGAFRT LFGGMSWITQ  750
2022_IIId  KNPADTGHGT VIVELQYTGS NGPCRVPISV TANLMDLTPV GRLVTVNPFI STGGANNKVM IEVEPPFGDS YIVVGRGTTQ INYHWHKEGS SIGKALATTW KGAQRLAVLG DTAWDFGSIG GVFNSIGKAV HQVFGGAFRT LFGGMSWITQ  750
2023_IIIa  KNPADTGHGT VIVELQYTGS NGPCRVPISV TANLMDLTPV GRLVTVNPFI STGGANNKVM IEVEPPFGDS YIVVGRGTTQ INYHWHKEGS SIGKALATTW KGAQRLAVLG DTAWDFGSIG GVFNSIGKAV HQVFGGAFRT LFGGMSWITQ  750
2023_IIId  KNPADTGHGT VIVELQYTGS NGPCRVPISV TANLMDLTPV GRLVTVNPFI STGGANNKVM IEVEPPFGDS YIVVGRGTTQ INYHWHKEGS SIGKALATTW KGAQRLAVLG DTAWDFGSIG GVFNSIGKAV HQVFGGAFRT LFGGMSWITQ  750

Envelope | NS1
2003_V     GLLGALLLWM GLQARDRSIS LTLLAVGGIL IFLATSVQAD SGCAIDLQRR ELKCGGGIFV YNDVEKWKSD YKYFPLTPTG LARVIQEAHA NGICGIRSTS RLEHLMWENI QRELNAIFED NEIDLSVVVQ EDPKHYRRAP QRLKKLEDEL  900
2016_IIId  GLLGALLLWM GLQARDRSIS LTLLAVGGIL IFLATSVQAD SGCAINLQRR ELKCGGGIFV YNDVEKWKSD YKYFPLTPTG LARVIQEAHA NGICGIRSTS RLEHLMWESI QKELNAIFED NEIDLSVVVQ EDPKYYKRAP RRLKKLEDEL  900
2017_IIIa  GLLGALLLWM GLQARDRSIS LTLLAVGGIL IFLATSVQAD SGCAIDLQRR ELKCGGGIFV YNDVEKWKSD YKYFPLTPTG LARVIQEAHA NGICGIRSTS RLEHLMWESI QKELNAIFED NEIDLSVVVQ EDPKYYKRAP RRLKKLEDEL  900
2022_IIId  GLLGALLLWM GLQARDRSIS LTLLAVGGIL IFLATSVQAD SGCAIDLQRR ELKCGGGIFV YNDVEKWKSD YKYFPLTPTG LARVIQEAHA NGICGIRSTS RLEHLMWESI QKELNAIFED NEIDLSVVVQ EDPKYYKRAP RRLKKLEDEL  900
2023_IIIa  GLLGALLLWM GLQARDRSIS LTLLAVGGIL IFLATSVQAD SGCAIDLQRR ELKCGGGIFV YNDVEKWKSD YKYFPLTPTG LARVIQEAHA NGICGIRSTS RLEHLMWESI QKELNAIFED NEIDLSVVVQ EDPKYYKRAP RRLKKLEDEL  900
2023_IIId  GLLGALLLWM GLQARDRSIS LTLLAVGGIL IFLATSVQAD SGCAIDLQRR ELKCGGGIFV YNDVEKWKSD YKYFPLTPTG LARVIQEAHA NGICGIRSTS RLEHLMWESI QKELNAIFED NEIDLSVVVQ EDPKYYKRAP RRLKKLEDEL  900

NS1
2003_V     DYGWKKWGKT LFMEPKLGNN TFVVDGPETK ECPTANRAWN SFKVEDFGFG MVFTRLWLTI REGNTTECDS AIIGTAIKGD RAVHSDLSYW IESKKNGTWQ LERAVMGEVK SCTWPETHTL WGDGVVESEM IIPVTLGGPK SHHNKRTGYH  1050
2016_IIId  NYGWKKWGKT LFMEPKLGNN TFVVDGPETK ECPTANRAWN SFKVEDFGFG MVFTRLWLTI REENTTECDS AIIGTAIKGD RAVHSDLSYW IESKKNGTWQ LERAVMGEVK SCTWPETHTL WGDGVVESEM IIPVTLGGPK SHHNKRTGYH  1050
2017_IIIa  NYGWKKWGKT LFMEPKLGNN TFVVDGPETK ECPTANRAWN SFKVEDFGFG MVFTRLWLTI REENTTECDS AIIGTAIKGD RAVHSDLSYW IESKKNGTWQ LERAVMGEVK SCTWPETHTL WGDGVVESEM IIPVTLGGPK SHHNKRTGYH  1050
2022_IIId  NYGWKKWGKT LFMEPKLGNN TFVVDGPETK ECPTANRAWN SFKVEDFGFG MVFTRLWLTI REENTTECDS AIIGTAIKGD RAVHSDLSYW IESKKNGTWQ LERAVMGEVK SCTWPETHTL WGDGVVESEM IIPVTLGGPK SHHNKRTGYH  1050
2023_IIIa  NYGWKKWGKT LFMEPKLGNN TFVVDGPETK ECPTANRAWN SFKVEDFGFG MVFTRLWLTI REENTTECDS AIIGTAIKGD RAVHSDLSYW IESKKNGTWQ LERAVMGEVK SCTWPETHTL WGDGVVESEM IIPVTLGGPK SHHNKRTGYH  1050
2023_IIId  NYGWKKWGKT LFMEPKLGNN TFVVDGPETK ECPTANRAWN SFKVEDFGFG MVFTRLWLTI REENTTECDS AIIGTAIKGD RAVHSDLSYW IESKKNGTWQ LERAVMGEVK SCTWPETHTL WGDGVVESEM IIPVTLGGPK SHHNKRTGYH  1050

NS1 | NS2A
2003_V     TQTKGPWSEG EITLDFDYCP GTTVTVTEHC GNRGASLRTT TASGKLVTDW CCRSCSLPPL RYTTKDGCWY GMEIRPVKEE EAKLVKSRVT AGVAGGMEPF QLGLLVAFIA TQEVLKRRWT GKLTLTSLAV CLALLIFGNL TYMDLVRYLV  1200
2016_IIId  TQTKGPWSEG EITLDFDYCP GTTVTVTEHC GNRGASLRTT TASGKLVTDW CCRSCSLPPL RYTTKDGCWY GMEIRPVKEE EAKLVKSRVT AGVAGGMEPF QLGLLVAFIA TQEVLKRRWT GKLTLTSLAV CLALLIFGNL TYMDLVRYLV  1200
2017_IIIa  TQTKGPWSEG EITLDFDYCP GTTVTVTEHC GNRGASLRTT TASGKLVTDW CCRSCSLPPL RYTTKDGCWY GMEIRPVKEE EAKLVKSRVT AGVAGGMEPF QLGLLVAFIA TQEVLKRRWT GKLTLTSLAV CLALLIFGNL TYMDLVRYLV  1200
2022_IIId  TQTKGPWSEG EITLDFDYCP GTTVTVTEHC GNRGASLRTT TASGKLVTDW CCRSCSLPPL RYTTKDGCWY GMEIRPVKEE EAKLVKSRVT AGVAGGMEPF QLGLLVAFIA TQEVLKRRWT GKLTLTSLAV CLALLIFGNL TYMDLVRYLV  1200
2023_IIIa  TQTKGPWSEG EITLDFDYCP GTTVTVIEHC GNRGASLRTT TASGKLVTDW CCRSCSLPPL RYTTKDGCWY GMEIRPVKEE EAKLVKSRVT AGVAGGMEPF QLGLLVAFIA TQEVLKRRWT GKLTLTSLAV CLALLIFGNL TYMDLVRYLV  1200
2023_IIId  TQTKGPWSEG EITLDFDYCP GTTVTVTEHC GNRGASLRTT TASGKLVTDW CCRSCSLPPL RYTTKDGCWY GMEIRPVKEE EAKLVKSRVT AGVAGGMEPF QLGLLVAFIA TQEVLKRRWT GKLTLTSLAV CLALLIFGNL TYMDLVRYLV  1200

NS2A
2003_V     LVGTAFAEMN TGGDVIHLAL VAVFKVQPAF LAGLFLRMQW SNQENILMVI GAAFLQMAAN DLKLEVLPIL NAMSIAWMLI RAMKEGKVAM CALPILCALT PGMRMAGLDV IRCLLLIIGI VTLLNERRES VAKKKGGYLL AAALCQAGLC  1350
2016_IIId  LVGTAFAEMN TGGDVIHLAL VAVFKVQPAF LAGLFFRMQW SNQENILMVV GAAFLQMAAN DLKLEVLPIL NAMSIAWMLI RAMKEGKVAM YALPILCALT PGMRMAGLDV IRCLLLIIGI VTLLNERRES VAKKKGGYLL AAALCQAGLC  1350
2017_IIIa  LVGTAFAEMN TGGDVIHLAL VAVFKVQPAF LAGLFFRMQW SNQENILMVI GAAFLQMAAN DLKLEVLPIL NAMSIAWMLI RAMKEGKVAM YALPILCALT PGMRMAGLDV IRCLLLIIGI VTLLNERRES VAKKKGGYLL AAALCQAGLC  1350
2022_IIId  LVGTAFAEMN TGGDVIHLAL VAVFKVQPAF LAGLFFRMQW SNQENILMVI GAAFLQMAAN DLKLEVLPIL NAMSIAWMLI RAMKEGKVAM YALPILCALT PGMRMAGLDV IRCLLLIIGI VTLLNERRES VAKKKGGYLL AAALCQAGLC  1350
2023_IIIa  LVGTAFAEMN TGGDVIHLAL VAVFKVQPAF LAGLFFRMQW SNQENILMVI GAAFLQMAAN DLKLEVLPIL NAMSIAWMLI RAMKEGKVAM YALPFLCALT PGMRMAGLDV IRCLLLIIGI VTLLNERRES VAKKKGGYLL AAALCQVGLC  1350
2023_IIId  LVGTAFAEMN TGGDVIHLAL VAVFKVQPAF LAGLFFRMQW SNQENILMVI GAAFLQMAAN DLKLEVLPIL NAMSIAWMLI RAMKEGKVAM YALPILCALT PGMRMAGLDV IRCLLLIIGI VTLLNERRES VAKKKGGYLL AAALCQAGLC  1350

NS2A | NS2B
2003_V     SPLIMMGGLI LAHPNGKRSW PASEVLTGVG LMCALAGGLL EFEETSMVVP FAIAGLMYIT YTVSGKAAEM WIEKAADITW EQNAEITGTS PRLDVDLDSH GNFKLLNDPG APVHLFALRF ILLGLSARFH WFIPFGVLGF WLLGKHSKRG  1500
2016_IIId  SPLIMMGGLI LAHPNGKRSW PASEVLTGVG LMCALAGGLL EFEETSMVVP FAIAGLMYIT YTVSGRAAEM WIEKAADITW EQNAEITGTS PRLDVDLDSH GNFKLLNDPG APVHLFALRF ILLGLSARFH WFIPFGVLGF WLLGKHSKRG  1500
2017_IIIa  SPLIMMGGLI LAHPNGKRSW PASEVLTGVG LMCALAGGLL EFEETSMVVP FAIAGLMYIT YTVSGRAAEM WIEKAADITW EQNAEITGTS PRLDVDLDSH GNFKLLNDPG APVHLFALRF ILLGLSARFH WFIPFGVLGF WLLGKHSKRG  1500
2022_IIId  SPLIMMGGLI LAHPNGKRSW PASEVLTGVG LMCALAGGLL EFEETSMVVP FAIAGLMYIT YTVSGRAAEM WIEKAADITW EQNAEITGTS PRLDVDLDSH GNFKLLNDPG APVHLFALRF ILLGLSARFH WFIPFGVLGF WLLGKHSKRG  1500
2023_IIIa  SPLIMMGGLI LAHPNGKRSW PASEVLTGVG LMCALAGGLL EFEETSMVVP FAIAGLMYIT YTVSGRAAEM WIEKAADITW EQNAEITGTS PRLDVDLDSH GNFKLLNDPG APVHLFALRF ILLGLSARFH WFIPFGVLGF WLLGKHSKRG  1500
2023_IIId  SPLIMMGGLI LAHPNGKRSW PASEVLTGVG LMCALAGGLL EFEETSMVVP FAIAGLMYIT YTVSGRAAEM WIEKAADITW EQNAEITGTS PRLDVDLDSH GNFKLLNDPG APVHLFALRF ILLGLSARFH WFIPFGVLGF WLLGKHSKRG  1500

NS3
2003_V     GALWDVPSPK VYPKCETKPG IYRIMTRGIL GTFQAGVGVM HEGVFHTMWH ATEGAVLRNG EGRLDPYAGD VRNDLISYGG PWKLSATWDG IEEVQMIAVA PGKPAINVQT TPGVFKTPFG TIGAVTLDFP KGTSGSPIIN KKGEIIGLYG  1650
2016_IIId  GALWDVPSPK VYPKCETKPG IYRIMTRGIL GTFQAGVGVM HEGVFHTMWH ATEGAVLRNG EGRLDPYAGD VRNDLISYGG PWKLSATWDG IEEVQMIAVA PGKPAINVQT TPGVFKTPFG TIGAVTLDFP KGTSGSPIIN KKGEIIGLYG  1650
2017_IIIa  GALWDVPSPK VYPKCETKPG IYRIMTRGIL GTFQAGVGVM HEGVFHTMWH ATEGAVLRNG EGRLDPYAGD VRNDLISYGG PWKLSATWDG IEEVQMIAVA PGKPAINVQT TPGVFKTPFG TIGAVTLDFP KGTSGSPIIN KKGEIIGLYG  1650
2022_IIId  GALWDVPSPK VYPKCETKPG IYRIMTRGIL GTFQAGVGVM HEGVFHTMWH ATEGAVLRNG EGRLDPYAGD VRNDLISYGG PWKLSATWDG IEEVQMIAVA PGKPAINVQT TPGVFKTPFG TIGAVTLDFP KGTSGSPIIN KKGEIIGLYG  1650
2023_IIIa  GALWDVPSPK VYPKCETKPG IYRIMTRGIL GTFQAGVGVM HEGVFHTMWH ATEGAVLRNG EGRLDPYAGD VRNDLISYGG PWKLSATWDG IEEVQMIAVA PGKPAINVQT TPGVFKTPFG TIGAVTLDFP KGTSGSPIIN KKGEIIGLYG  1650
2023_IIId  GALWDVPSPK VYPKCETKPG IYRIMTRGIL GTFQAGVGVM HEGVFHTMWH ATEGAVLRNG EGRLDPYAGD VRNDLISYGG PWKLSATWDG IEEVQMIAVA PGKPAINVQT TPGVFKTPFG TIGAVTLDFP KGTSGSPIIN KKGEIIGLYG  1650

NS3
2003_V     NGVLIGQGEY VSGIIQGERT EEPIPDAYNE EMLRKRKLTV LELHPGAGKT RKVLPQIIKD CIQKRLRTAV LAPTRVVACE IAEALKGLPI RYLTPAVKNE HQGNEIVDVM CHATLTQKLL TPTRVPNYQV YIMDEAHFID PASIAARGYI  1800
2016_IIId  NGVLIGQGEY VSGIIQGERT EEPIPDAYNE EMLRKRKLTV LELHPGAGKT RKVLPQIIKD CIQKRLRTAV LAPTRVVACE IAEALKGLPI RYLTPAVKNE HQGNEIVDVM CHATLTQKLL TPTRVPNYQV YIMDEAHFID PASIAARGYI  1800
2017_IIIa  NGVLIGQGEY VSGIIQGERT EEPIPDAYNE EMLRKRKLTV LELHPGAGKT RKVLPQIIKD CIQKRLRTAV LAPTRVVACE IAEALKGLPI RYLTPAVKNE HQGNEIVDVM CHATLTQKLL TPTRVPNYQV YIMDEAHFID PASIAARGYI  1800
2022_IIId  NGVLIGQGEY VSGIIQGERT EEPIPDAYNE EMLRKRKLTV LELHPGAGKT RKVLPQIIKD CIQKRLRTAV LAPTRVVACE IAEALKGLPI RYLTPAVKNE HQGNEIVDVM CHATLTQKLL TPTRVPNYQV YIMDEAHFID PASIAARGYI  1800
2023_IIIa  NGVLIGQGEY VSGIIQGERT EEPIPDAYNE EMLRKRKLTV LELHPGAGKT RKVLPQIIKD CIQKRLRTAV LAPTRVVACE IAEALKGLPI RYLTPAVKNE HQGNEIVDVM CHATLTQKLL TPTRVPNYQV YIMDEAHFID PASIAARGYI  1800
2023_IIId  NGVLIGQGEY VSGIIQGERT EEPIPDAYNE EMLRKRKLTV LELHPGAGKT RKVLPQIIKD CIQKRLRTAV LAPTRVVACE IAEALKGLPI RYLTPAVKNE HQGNEIVDVM CHATLTQKLL TPTRVPNYQV YIMDEAHFID PASIAARGYI  1800
```

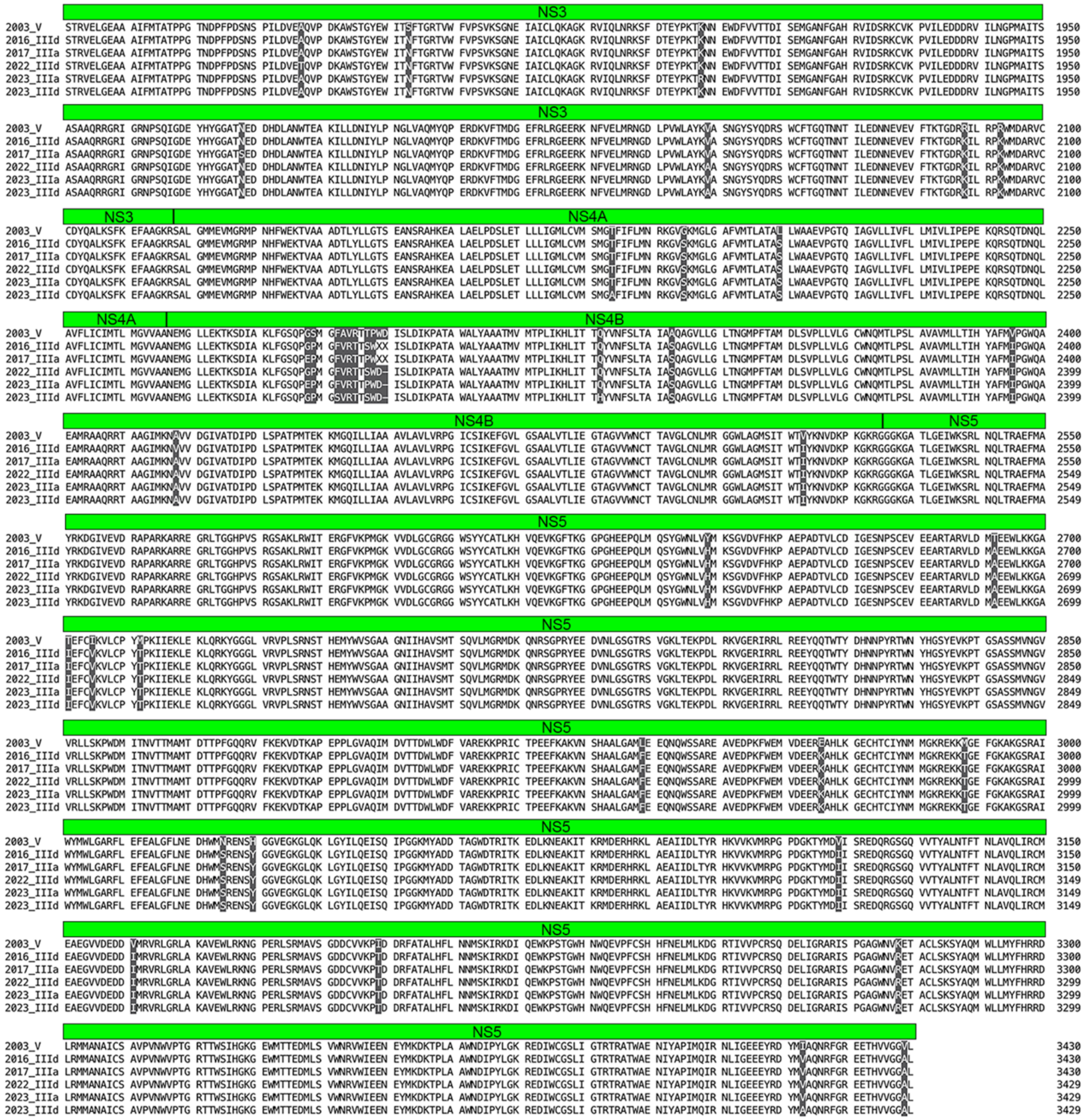
NS3
NS4A
NS4B
NS5

**Supplemental Figure 3**: Genome length amino acid alignment of six SLEV strains used in these studies. NS is non-structural. Dark shading shows locations of variant amino acids.
